# Predicting MammaPrint Recurrence Risk from Breast Cancer Pathological Images Using a Weakly Supervised Transformer

**DOI:** 10.1101/2025.04.26.648504

**Authors:** Chaoyang Yan, Linwei Li, Xiaolong Qian, Yang Ou, Zhidong Huang, Zhihan Ruan, Wenting Xiang, Hong Liu, Jian Liu

## Abstract

Recurrence related to poor prognosis is a leading cause of mortality in patients with breast cancer (BC). The MammaPrint (MP) genomic assay is designed to stratify recurrence risk and evaluate chemotherapy benefits for early-stage HR+/HER2- BC patients. However, MP fails to reveal spatial tumor morphology and is limited by high costs. In this study, we establish a BC MP cohort and develop CPMP, a weakly supervised agent-attention transformer model, to predict MP recurrence risk from annotation-free BC histopathological slides. CPMP achieves an AUROC of 0.824 ± 0.03 in predicting MP risk groups. We further leveraged CPMP for spatial and morphological analyses to explore histological patterns associated with MP risk groups. The model reveals tumor spatial localization at the whole-slide level and highlights distinct intercellular interaction patterns of MP groups. It also characterizes the diversity in tumor morphology and uncovers MP High-specific, Low-specific, and colocalized morphological phenotypes that differ in quantitative cellular composition. Prognostic evaluation in the external cohort exhibits significant stratification of distant metastasis risk (HR: 3.14, *p*-value = 0.0014), underscoring the prognostic power of CPMP. These findings demonstrate the capability of CPMP in MP risk prediction, offering a flexible supplement to genomic risk assessment in early-stage BC.

## 1. Introduction

Breast cancer (BC) is the most common cancer and the leading cause of cancer deaths among women worldwide^[1]^. Recurrence and distant metastasis are significant factors that contribute to poor prognostic outcomes and are the primary causes of death among BC patients^[2,3]^. Recurrence risk stratification is essential for informing therapeutic regimens in the clinical management of BC. Currently, recurrence and metastasis prediction remains a challenging task, typically assessed through clinical characteristics, the Nottingham Grading system, or the Tumor-Node-Metastasis (TNM) staging system^[4]^. However, it can be challenging to determine the optimal treatment for individuals using these schemes, which may lead to overtreatment in some cases.

The 70-gene MammaPrint^®^ (MP) test, which has drawn attention among clinicians, is specifically designed to evaluate recurrence risk in early-stage Hormone Receptor positive (HR+), Human Epidermal Growth Factor Receptor 2 negative (HER2-) invasive BC patients using formalin-fixed paraffin-embedded (FFPE) or fresh frozen tissue^[5–7]^. This test estimates the 10-year risk of distant metastasis and classifies patients into Low or High-risk groups for recurrence. The MP test effectively identifies High-risk patients who may benefit from adjuvant chemotherapy and Low-risk patients who, despite having clinical high-risk features, may safely be exempt from chemotherapy^[7,8]^. Although the genomic MP test offers significant clinical value, it falls short in revealing the spatial characteristics of tumors for recurrence risk diagnosis. Additionally, the MP test is time-consuming, labor-intensive, and requires substantial resources. Its high cost presents a financial barrier to many patients^[9]^. This situation ultimately hinders patients from receiving precise treatments based on a risk assessment of their condition. Therefore, there is a need to develop an accurate and cost-efficient method for patient recurrence risk stratification.

Histopathology provides extensive insights into the spatial distribution of tissue, cellular interactions, and tumor morphology^[10]^, establishing itself as the “gold standard” for cancer diagnosis. The increasing accessibility of digital histopathological whole-slide images (WSIs), coupled with advancements in artificial intelligence (AI) technology, has propelled the research of computational pathology (CPATH) forward^[11–13]^ and made remarkable strides in molecular biomarker discovery^[14–20]^- Recent studies^[21–25]^ have demonstrated the potential of deep learning to extract prognostic information from BC histopathology. Wahab et al. developed an AI-based BRACE marker from stromal and tumor heterogeneity to predict survival in early-stage luminal BC patients using routine WSIs^[26]^. Chen et al. created a computational pathology framework that enhances risk stratification for early-stage BC patients within the risk categories defined by the Oncotype DX^®^ (ODX) assay^[27]^. Furthermore, Howard et al. integrated clinicopathological variables with image-based features to predict recurrence risk scores based on ODX assay results^[28]^. These studies underscore the value of identifying morphological biomarkers from pathological WSIs to complement genomic assays in enhancing prognostic stratification. However, the application of deep learning to predict recurrence risk based on paired MP genomic data and histopathological images remains unexplored. There is a critical gap in research that leverages the MP test’s genomic foundation to characterize histological patterns relevant to distant metastasis risk stratification and patient prognosis.

To bridge this gap, we construct the clinical MP cohort of HR+/HER2- invasive BC patients who underwent standardized MP genomic testing at Tianjin Medical University Cancer Institute and Hospital (TJMUCH), National Clinical Research Center for Cancer, China. We present an AI-driven computational pathology framework (CPMP) for MP- informed recurrence risk assessment using histopathological WSIs from early-stage BC patients. We develop a spatially-aware, weakly supervised learning model with a regression-based design and an agent-attention transformer paradigm to predict continuous recurrence risk scores from annotation-free digital WSIs. We further generate spatial attention heatmap visualization at the whole-slide level to reveal intratumoral spatial localization, as well as the cellular interaction patterns associated with genomic MP within the tumor ecosystem. We characterize the phenotypic diversity in tumors and uncover High-specific, Low-specific, and colocalized morphological phenotypes among MP risk groups, each differing in their quantitative cellular compositions. Moreover, we expand our model to conduct the generalization analysis in the external TCGA-BRCA cohort and demonstrate its predictive capability for risk stratification across multiple prognostic indicators. The CPMP framework is capable of predicting MP recurrence risk, facilitating the discovery of prognostic knowledge, and has the potential to serve as a flexible complement to the standard clinical diagnostic workflow for early-stage BC patients. Here, ’flexible’ refers to cost-efficient, rapid recurrence risk assessment directly from H&E-stained slides, while also providing spatial and morphological information that enhances the genomic MP test.

## 2. Results

### 2.1. Overview of the CPMP model and study design

We establish an early-stage HR+/HER2- invasive BC cohort, TJMUCH-MP, comprising 477 female patients (Figure S1 and S2a). The patient cohort is composed of clinicopathologic characteristics, digital WSIs, and genomic MP diagnostic results (Figure S1b-d). The clinicopathological characteristics of patients in the TJMUCH-MP cohort are presented in Table S1. MP diagnostic tests have been performed on tumor specimens of patients who met the inclusion criteria with invasive BC, HR+/HER2-, staged T1-2, lymph node-negative or 1- 3 metastases from the StGallen recommendations^[29]^. Patients are categorized into High-risk and Low-risk based on the risk values of the genomic MP test (Figure S1c and S2b).

We present CPMP, a weakly supervised multiple instance learning (MIL) model for MP- informed recurrence risk assessment of BC patients based on annotation-free histopathological WSIs (**Figure 1**a). Our proposed CPMP framework first preprocesses WSIs for the tissue region identification and tessellates local tiles with a size of 256×256 within the foreground regions at 20× objective magnification (Figure S2c). The median number of tiles across WSIs in the TJMUCH-MP cohort is 8,229, and the distribution of the number of tiles in WSIs has been reported (Figure S2d). Then, the CPMP framework captures diverse patterns and generates a low-dimensional feature representation of local tile instances using self-supervised foundation models (Figure S2e). Given that the spatial relationships among tissue regions suggest cellular interactions that potentially reveal biological behaviors and prognostic significance, the Positional coordinate information Embedding (PE) of each tile is encoded and integrated with low-dimensional tissue embeddings to enhance spatial correlation among local regions. CPMP introduces the transformer architecture (Figure 1b) for feature aggregation and employs the agent-attention mechanism^[30]^ (Figure 1c) to achieve a balance between global representation capability and computational efficiency. Additionally, CPMP utilizes a regression-based multi-layer perceptron (MLP) predictor to generate a continuous numerical value representing genomic MP risk scores.

**Figure 1.**
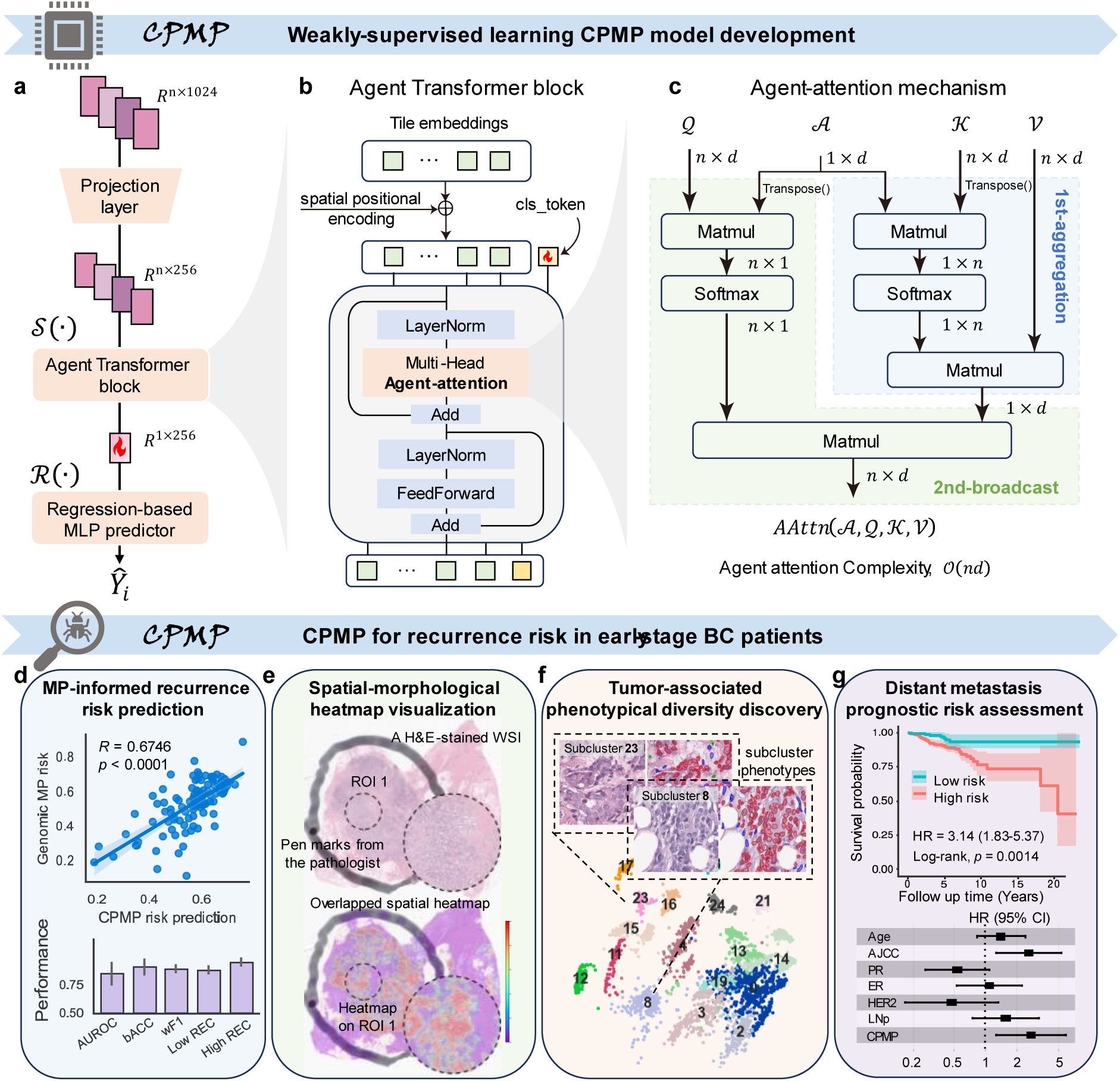
Overview of CPMP and study design. **a.** Depiction of the proposed weakly-supervised transformer architecture CPMP based on a multi-instance learning paradigm. It first performs linear projection on the feature embeddings of local tiles. Then it employs the Agent Transformer architecture for feature aggregation to obtain a slide-level global feature representation, followed by a regression-based MLP predictor head that predicts the slide-level results for the case. **b.** The Agent Transformer block adopted in the CPMP model. We introduced a 2-dimensional absolute coordinate encoding method to incorporate the spatial positional information and added the resulting positional embeddings to the projected tile embeddings. Then we initialized a class token (cls_token) and concatenated it with the spatial-aware tile embeddings. This Agent Transformer block consists of 1 layer with four heads for global information aggregation. **c.** The Agent-attention mechanism utilized in the Agent Transformer block. The Agent-attention paradigm consists of a stacked state of two Softmax attention operations, comprising the first phase of 1st-aggregation (colored in light blue) and the second phase of 2nd-broadcast (colored in light green). **d-g.** Flowchart of the study design: (**d**) MP-informed recurrence risk prediction, (**e**) Spatial-morphological attention heatmap visualization, (**f**) Tumor-associated phenotypical diversity discovery, and (**g**) Distant metastasis prognostic risk assessment. These also summarize the clinical value of CPMP. MLP = Multi-Layer Perceptron, MP = MammaPrint.

We developed and trained the CPMP model to predict MP-informed patient-level recurrence risk (Figure 1d) using the clinical TJMUCH-MP cohort. In addition, we leverage CPMP to generate whole-slide-level spatial attention heatmaps using histopathological slides (Figure 1e), which provide key morphological patterns and spatial localization associated with MP recurrence risk. We investigate tumor morphological patterns by clustering the learned tile embeddings from CPMP to characterize tumor diversity and identify specific, colocalized morphological phenotypes across MP risk groups (Figure 1f). Moreover, we generalize our trained CPMP model to the external TCGA-BRCA cohort for recurrence risk stratification and prognostic analysis, aiming to validate the potential capabilities of CPMP in assessing distant metastasis risk (Figure 1g). More details are provided in the Methods section.

### 2.2. CPMP outperforms other methods for MP recurrence risk prediction

The model’s ability to predict the recurrence risk from histopathological WSIs was measured on both continuous risk scores and discrete risk groups using the TJMUCH-MP cohort. To mitigate data bias and underlying scanning effects (Figure S2f), we employed a patient-level 5-Time 5-Fold cross-validation strategy (Figure S3a) for model development and evaluation. Our model achieved an average Spearman correlation coefficient of 0.648 [*p* < 0.0001, range: 0.606-0.684], indicating a favorable consistency between genomic MP risk scores and recurrence risk prediction probabilities (**Figure 2**a and S3d-i). In classifying the MP recurrence risk groups, CPMP attained an average area under the receiver operating characteristic curve (AUROC) of 0.824 ± 0.03 [range: 0.789-0.853] (Figure 2b), with a balanced accuracy of 0.772±0.03 [range: 0.750-0.773]. CPMP provides Low-as-positive and High-as-positive modes, which means that we aim to focus on Low-risk patients to avoid overtreatment, or, conversely, to monitor High-risk patients who may be at risk of recurrence. CPMP reached average areas under the precision-recall curves (AUPRCs) of 0.873±0.02 [range: 0.835-0.900] on the Low-as-positive mode (Figure S3b) and 0.774±0.01 [range: 0.751-0.795] on the High-as-positive mode (Figure S3c).

**Figure 2.**
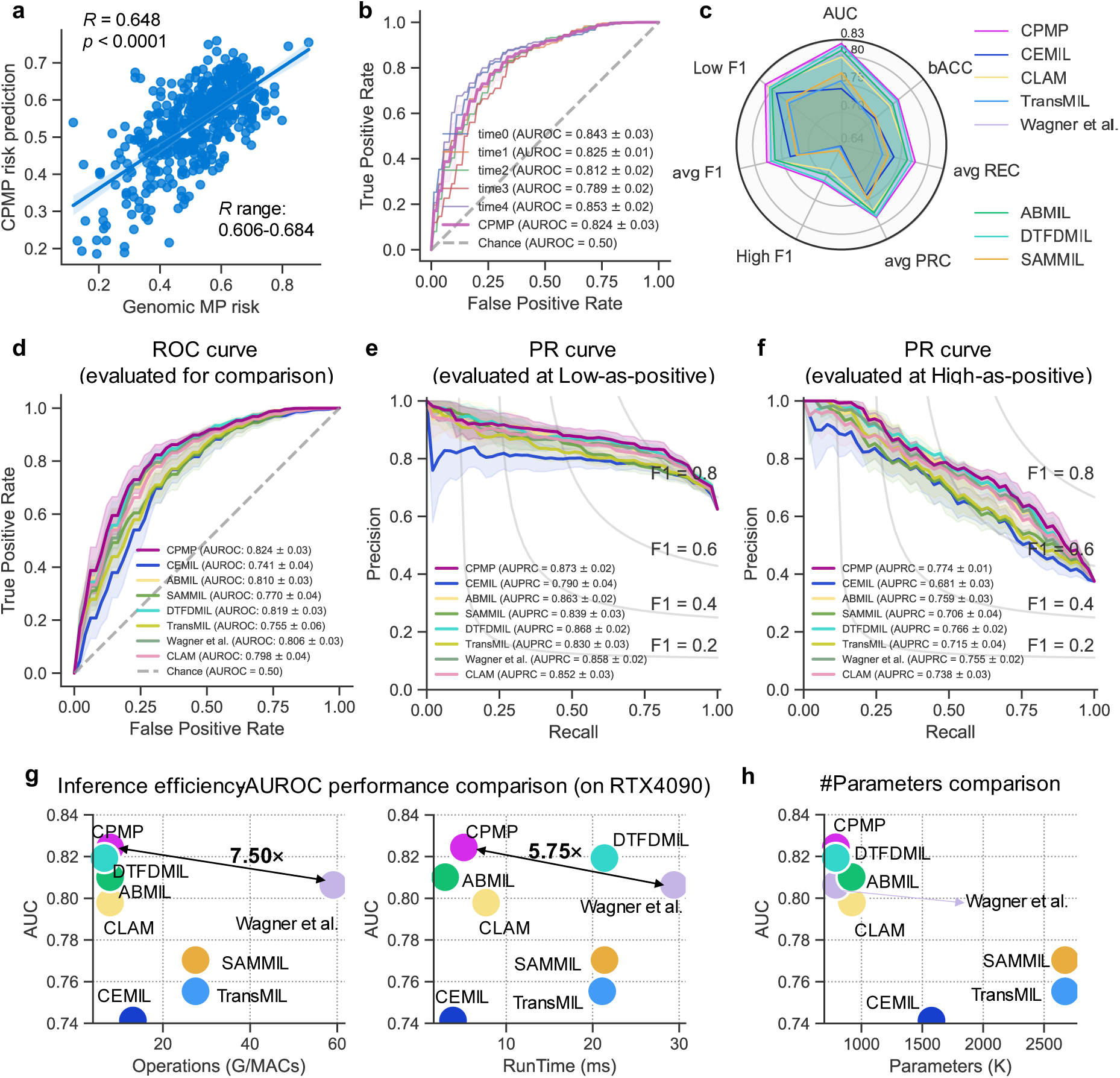
Performance evaluation and comparison of CPMP with state-of-the-art methods for MP recurrence risk prediction. **a**. Scatter plot showing the correlation between the continuous genomic MP risk score and the continuous CPMP-predicted risk probability for each patient in the test set, with a Spearman correlation coefficient (R) of 0.648 (*p*-value < 0.0001). The fitted linear regression line is also shown. **b**. ROC curve of the CPMP model for MP recurrence risk group prediction across 5-Time experiments (time0 to time4). **c**. Radar chart comparing the overall performance of CPMP with other comparative models (CEMIL, CLAM, TransMIL, Wagner et al., ABMIL, DTFDMIL, SAMMIL) across multiple metrics, including AUROC, balanced accuracy (bACC), average F1 score, average recall (avg REC), and average precision-recall curve (avg PRC). **d-f**. Comprehensive comparison of CPMP with seven state-of-the-art weakly supervised MIL methods using multiple metrics: (**d**) ROC curves, (**e**) Precision-Recall (PR) curves at the Low-as-positive mode, and (**f**) PR curves at the High-as-positive mode. Low-as-positive is the condition that we aim to focus on Low-risk patients to avoid overtreatment. High-as-positive is to observe the High-risk patients who may be at risk of recurrence. **g**. Evaluation of the efficiency-AUC performance trade-off of CPMP and comparative models in terms of AUC vs. computational operations (G/MACs) and AUC vs. inference runtime (ms). The data points are colored by the corresponding method. The results were obtained on a workstation equipped with two RTX 4090 GPUs. **h**. Evaluation of the number of parameters (K) of trained models for CPMP and comparative models. ROC=Receiver Operating Characteristic, PR=Precision-Recall, AUC=Area Under the Receiver Operating Characteristic.

The performance of our spatially aware, agent-attention transformer model was further compared with other weakly supervised MIL methods, including CEMIL^[31]^, SAMMIL^[32]^, DTFDMIL^[33]^, ABMIL^[34]^, CLAM^[35]^, Wagner et al.^[36]^, and TransMIL^[37]^. Details of all comparative models are provided in the Methods section. CPMP outperformed all comparative approaches in MP recurrence risk prediction in terms of systematic metrics, including AUROC, balanced accuracy, weighted average recall, precision, F1 scores, ROC curves, and precision-recall curves (Figure 2c-f and Table S2). The average AUROC value of the CPMP model is 0.824, while those of other methods, namely CEMIL, SAMMIL, DTFDMIL, CLAM, TransMIL, ABMIL, and Wagner et al., are 0.741±0.04, 0.770±0.04, 0.819 ± 0.03, 0.798 ± 0.04, 0.755 ± 0.06, 0.810 ± 0.03, and 0.806 ± 0.03, respectively. It is noteworthy that the Transformer-based methods (Wagner et al. and TransMIL) obtained acceptable performance but fell short in comparison with our transformer-based CPMP. We visualized the predicted risk probabilities from comparative methods alongside genomic MP risk scores of patients (Figure S4a-h). The results indicate that these classification-based methods tend to predict risk probabilities that are skewed toward both ends. Meanwhile, CPMP achieved performance improvements ranging from 4.8% (compared to DTFDMIL) to 33.6% (compared to CEMIL) in the Spearman correlation coefficient over the comparative methods (Figure S4a, b, d). This is attributed to the regression-based design in our method. The computational operations and runtime costs for inference were conducted on a workstation with RTX 4090 GPUs for comparison. We can see that CPMP reduces inference operations (G/MACs) by 7.50× times and increases inference time by 5.75× times, while improving AUC performance by 2.2% (*p* < 0.05, statistically significant difference), compared with the Wagner et al. transformer (Figure 2g). Also, CPMP requires fewer parameters than other methods (Figure 2h). CPMP attains the most superior efficiency-AUC performance during inference, striking a balance between slide representation capability and computational complexity.

### 2.3. CPMP highlights its effectiveness in leveraging annotation-free WSIs

Blurry areas, pen markings, and non-tissue regions are inevitably present in annotation-free histopathological WSIs from real-world clinical data, introducing noise into the training data. Thus, we investigated CPMP’s sensitivity to noise by removing noisy tiles. Specifically, we performed zero-shot classification to determine tissue types (tumor, adipose, stroma, immune infiltrates, gland, necrosis or hemorrhage, background or black, and non) across all tiles through the visual-language foundation model PLIP^[38]^ (Figure S5a, b). Local tile images along with their corresponding zero-shot classification types were visualized (Figure S5c). Tiles classified as “background or black” and “non” types are typically blurry, pen-marked, or lacking morphological detail. In contrast, tiles identified as the six specific tissue types (tumor, adipose, stroma, immune infiltrates, gland, and necrosis or hemorrhage) contain rich morphological information and are thus utilized for training the other model. Evaluation results for both models indicate that the model with noisy tile filtering achieved comparable performance to the CPMP model without such filtering (**Figure 3**a and Table S3), showing no statistically significant difference between the two. This reflects the robustness of our weakly supervised CPMP approach in handling local noisy information within annotation-free slides, making it more suitable for clinical scenarios.

**Figure 3.**
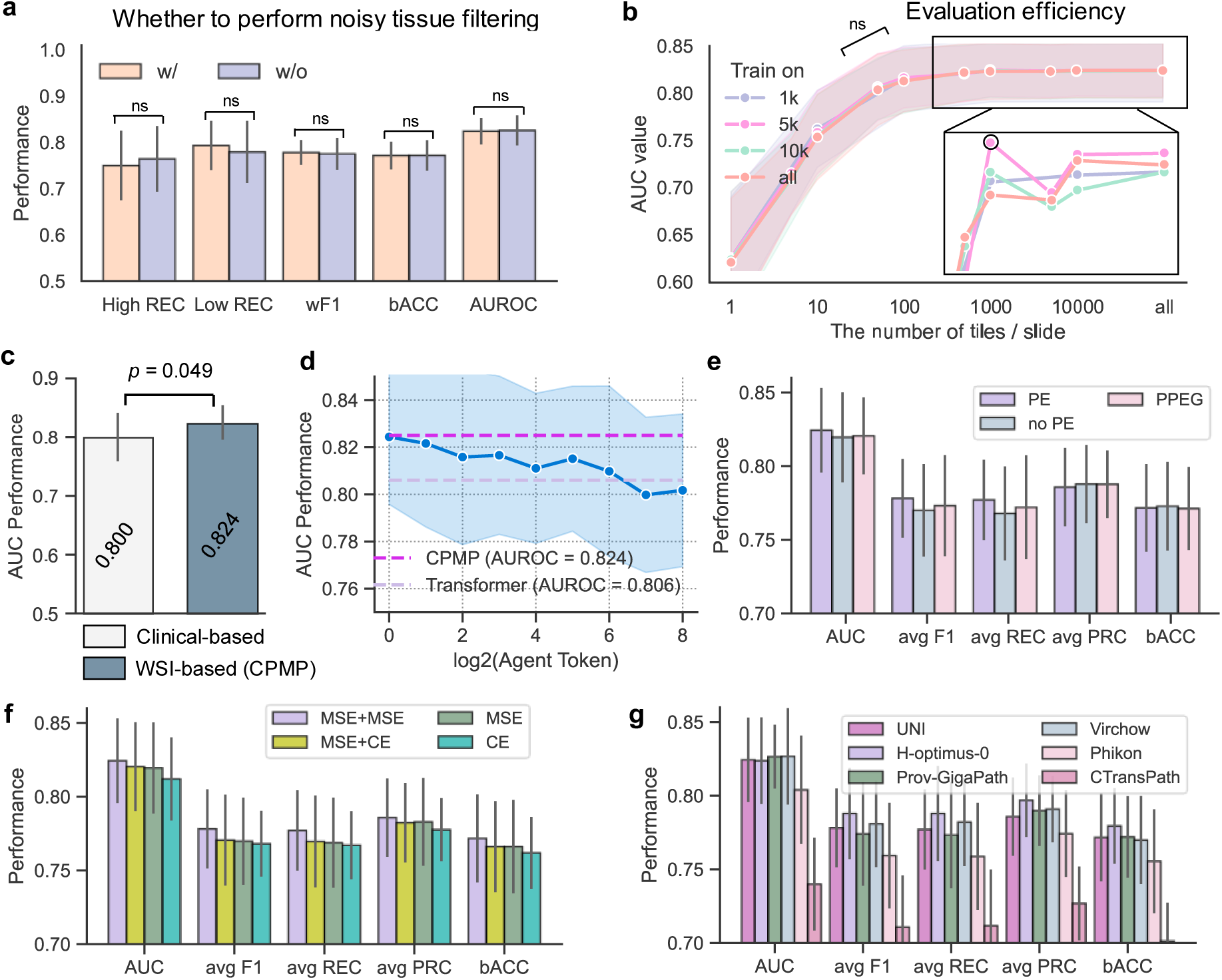
Effectiveness overview of CPMP in leveraging annotation-free WSIs. **a.** Comparison of the performance of CPMP and the model with noisy tissue-type filtering in terms of 5 evaluation metrics. The Wilcoxon signed-rank test analysis was performed. **b.** AUROC values of models trained on different sets with a specific number of tiles randomly sampled from each slide for evaluation. The AUROC values were averaged across 5-Time experiments. **c.** Comparison of the AUROC performance of CPMP and the clinical-based logistic regression model trained on clinicopathological features only. The Wilcoxon signed-rank test analysis was performed. **d.** AUROC performance trend evaluation of the agent- transformer models with a varying number of agent tokens. The light blue shading represents the standard deviation interval of performance across 5-Time 5-Fold experiments, while the dashes are the AUROC performance lines of CPMP and the self-attention transformer. **e.** Performance evaluation of the experimental models equipped with different positional encoding configurations, including PE, No-PE, and PPEG. **f.** Performance evaluation of the experimental models equipped with different loss functions. CE: Classification-based Cross-Entropy loss, MSE: Regression-based Mean-Square-Error loss. MSE+CE: Regression-based MSE loss and CE loss with soft label, MSE+MSE: Regression-based dual MSE loss strategy. **g.** Performance evaluation of the experimental models equipped with different feature extraction foundation models, including UNI, Virchow, H-optimus-0, Phikon, Prov-GigaPath, and CTransPath. w/=With, w/o=Without, ns=Statistically no significant difference, AUC=Area Under the receiver operating Characteristic, PE=Positional coordinate information Embedding, PPEG=Pyramid Position Encoding Generator, CE=Cross-Entropy, MSE=Mean Square Error.

We next evaluated the scale flexibility of CPMP in the number of local tiles in WSIs. We conducted random sampling from the set of local tiles in each WSI, employing sample sizes of 1,000, 5,000, and 10,000 tiles as subsets for model training. The trained models on various subsets of tiles exhibit consistent performance trends on various numbers of tiles, with differences in AUROC remaining within 0.01 compared to the model using all tiles (*p*-value > 0.05, indicating no statistically significant differences, Figure 3b and Table S4). Notably, these models demonstrated robust performance with AUROC values exceeding 0.810 when making predictions based on as few as 100 tiles up to the full set of tiles. This suggests that CPMP is both scalable and efficient in recurrence risk inference using histopathological WSIs. We further compared CPMP with a clinical-based logistic regression model that was trained on clinicopathological features for MP recurrence risk prediction for patients. The detailed settings for this clinical-based model are listed in the Methods section and Table S5. Clinical-based model obtained an acceptable AUC value of 0.800 (Figure 3c and Table S6), but it was relatively inferior to CPMP, especially given the fact that the clinical-based logistic regression model requires molecular information such as hormone receptor status and Ki-67 percentages, whereas CPMP relies solely on routine H&E-stained histopathological slides.

We conducted ablation studies on CPMP with respect to attention mechanisms, spatial positional encoding schemes, predictor head strategies, and tile-level feature representation models. The design outlines and the experimental configurations utilized for the ablation models are illustrated in Figure S6. Details about these modules are provided in the Methods section. The quantitative results reported that our CPMP method with the agent-attention mechanism brought a 1.8% improvement in AUROC over the self-attention transformer model, and it was insensitive to the number of agent tokens (Figure 3d and Table S7). Our PE-based model outperformed the PPEG-based^[37]^ and No-PE models in AUROC, weighted average recall, and F1 score (Figure 3e and Table S8). Additionally, a similar improvement can be observed in our regression-based mean square error supervision scheme when compared to the cross-entropy approach (Figure 3f and Table S9). In the ablation study of tile-level feature representation, the Prov-GigaPath^[39]^ and Virchow^[40]^ models excelled in UNI^[41]^ in terms of AUROC (0.827 vs. 0.824), while H-optimus-0^[42]^ achieved the highest values across other metrics (Figure 3g and Table S10). Additionally, the performance of CTransPath^[43]^ and Phikon^[44]^ was inferior to that of other models in representing tile embeddings for MP recurrence risk prediction in the TJMUCH-MP cohort. Given that the embedding dimensions of tile representations of Prov-GigaPath, Virchow, and H-optimus-0 are larger than those of UNI, and considering the balance with computational efficiency, we selected the UNI-based model for the subsequent analyses reported in this study.

### 2.4. CPMP reveals the spatial localization of tumors at the whole-slide level

Our proposed weakly supervised CPMP model not only predicts patient recurrence risk using spatial histopathological WSIs but also infers tile-level attention scores. These scores can offer insights into spatial factors that influence patient outcomes from the perspective of the whole-slide level. We performed whole-slide-level attention heatmap inference on test cases. Spatial heatmap results were generated by mapping the attention scores of tiles to their corresponding locations within the slide. This allows us to visualize the contributions of different regions and display key morphological patterns. Details are provided in the Methods section. The representative histopathological WSIs selected from genomic MP Low-risk and High-risk groups were presented, along with their spatial visualization results at both the whole-slide and region-of-interest (ROI) levels (**Figure 4**a-c, e-g and S7a-c, e-g). Genomic MP diagnostic results and recurrence risk values predicted by the CPMP model were also marked in the box (Figure 4a, e, and S7a, e). One can observe that CPMP exhibits varying levels of attention confidence in tissue regions, with high confidence indicated in red and low confidence in blue-purple, at the whole-slide level for both Low-risk and High-risk cases (Figure 4b, f and S7b, f). The pathologists confirmed that the areas identified as high confidence closely correspond to regions where tumor cells proliferate, based on their spatial locations. It can also be observed from the magnified ROIs that the high-attention heatmap focuses on the tumor regions and effectively outlines the boundary between tumor and normal stroma tissue (Figure 4c, g and S7c, g). Hot regions, as rendered by the spatial heatmap visualization, are highly consistent with the tumor regions pre-marked by experts (Figure S7e, f). CPMP further identifies the top 10 local tiles with the highest attention scores from each slide (Figure 4d, h, and S7d, h). These selected tiles display clear patterns of tumor morphology applicable to both Low-risk and High-risk cases. To quantitatively validate the interpretability of CPMP, breast pathologists manually annotated tumor regions on WSIs, allowing us to quantify the concordance between CPMP-generated high-attention regions (threshold=0.5) and pathologist-labeled ground-truth tumor regions (Figure S8 and S9). Multiple evaluation metrics confirmed the agreement: mean AUC=0.946, recall ratio=0.630±0.171, and Dice coefficient=0.622±0.126 (Table S11). These results demonstrate a desired alignment between model attention and expert annotations. A mean overlap ratio of 0.704 further indicates that the model’s attention is mainly focused on regions marked by pathologists as tumor tissue, providing quantitative support for the reliability of CPMP attention maps. The spatial heatmap results indicate that our model can comprehensively pinpoint tumor growth locations and capture the spatial localization of tumor ROIs at the whole-slide level. This capability of tumor spatial localization offers a visually explainable approach to recurrence risk prediction that complements the genomic MP test with valuable spatial insights.

**Figure 4.**
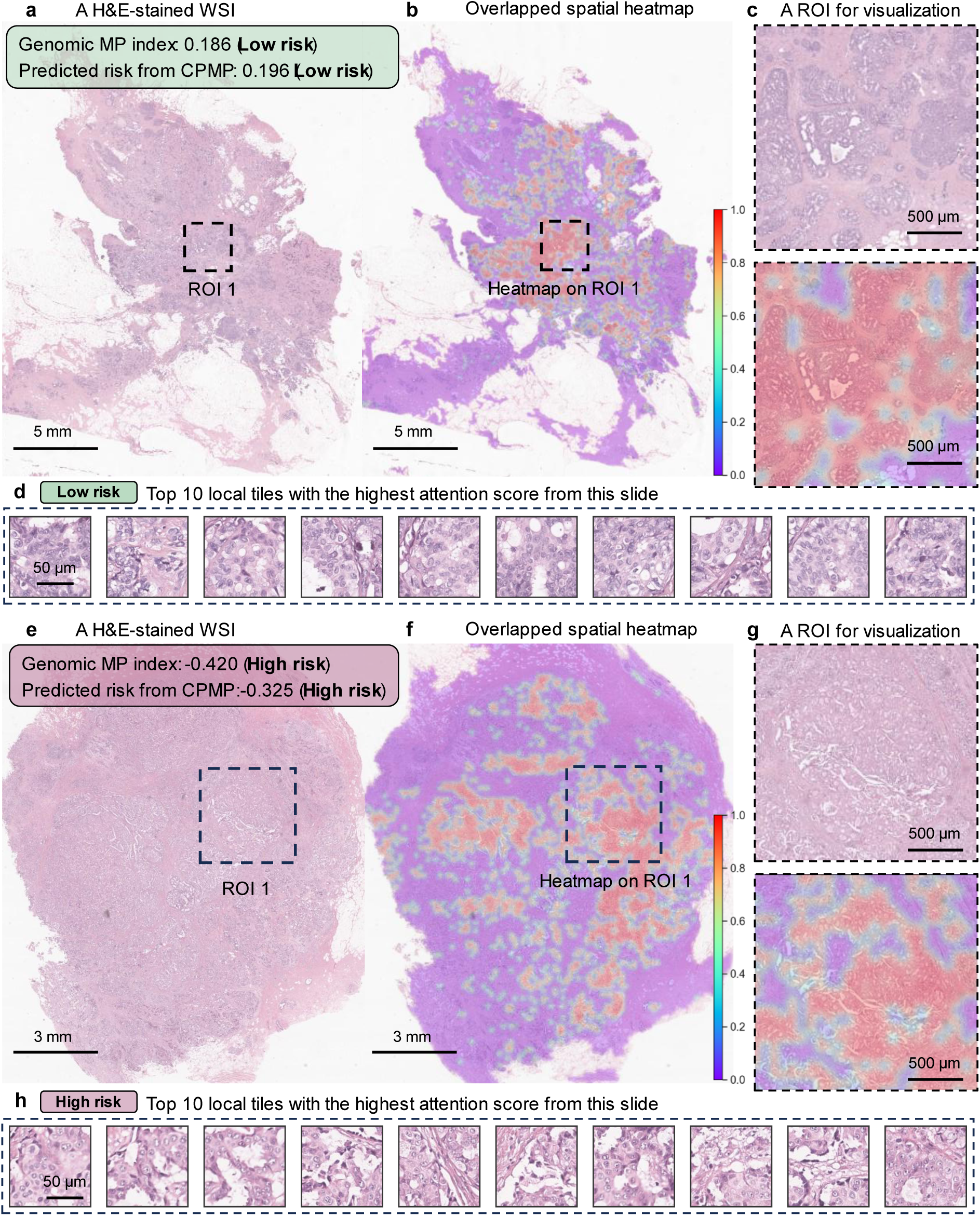
Heatmap visualization from CPMP revealing the spatial localization of tumors at the whole-slide level. a, **e.** Example H&E-stained WSIs of patients with genomic MP Low-risk (**a**) and High-risk (**e**) in the TJMUCH-MP cohort. Genomic MP diagnostic risk index and corresponding risk prediction results from CPMP are marked in the box. Scale bar: 5 mm / 3 mm. **b, f.** The overlaid spatial heatmap visualization for the corresponding Low-risk (**b**) and High-risk (**f**) cases at the whole-slide level. The tile attention scores that contributed to the final patient-level risk prediction were yielded from the agent-attention matrix of local tiles using the attention rollout method. Spatial heatmap results were then generated by mapping the attention scores of all local tiles to their corresponding locations within the WSIs, and they were superposed onto the histopathological WSIs for interpretability. Please see the Methods section. A colorbar was added on the lower right of the heatmap. Scale bar: 5 mm / 3 mm. **c**, **g.** Fine-grained pathological ROIs extracted from the WSIs and the corresponding spatial heatmap visualization results. Scale bar: 500 μm. **d, h.** Top 10 local tiles with the highest attention score from each slide for both MP Low-risk (**d**) and High-risk (**h**) cases. Scale bar: 50 μm. WSI=Whole Slide Image, ROI=Region of Interest, MP=MammaPrint diagnostic test.

### 2.5. CPMP deciphers cellular interaction patterns within the tumor ecosystem

To gain deeper insights into the intercellular spatial relationships within the tumor ecosystem across MP recurrence risk groups, high-attention ROIs localized by CPMP were extracted for simultaneous nuclear instance segmentation and classification (Figure S10a). HoverNet^[45]^ was utilized for the identification of five types of cell nuclei: neoplastic (tumor) cells, inflammatory cells, necrotic (dead) cells, connective (stroma) cells, and non-neoplastic epithelial cells. The contours of the identified nuclei were delineated on the fine-grained histopathological ROIs. From the nuclei segmentation visualization results for both Low-risk and High-risk ROIs (**Figure 5**a, b), we can observe that the identified cell nuclei are predominantly tumor cells (outlined in red), while stromal cells (indicated in blue) are sparsely distributed around the tumor tissue. The results of nuclei identification and visualization of the top 10 local tiles further support this finding (Figure 5c, e, and S10c-h). It is noteworthy that High-risk cases exhibit more dispersed spatial distribution among tumor cells compared to Low-risk cases. We thus quantified the cellular composition of the five cell types in each local tile and defined the density of tumor cells (TcD) as the number of identified neoplastic cells per square millimeter (Figure 5d, f). We can see that there is a clear difference in tumor cell density between the two MP risk groups. The number of neoplastic cells in Low-risk tiles is generally larger than that in High-risk tiles (median: 103 vs. 53, Low-risk vs. High-risk). The measured TcD values for Low-risk tiles are consistently greater than 4,000 tumor cells per mm²(median: 5,698, [range: 4,149-7,081]), while those for High-risk tiles are all below 4,000 neoplastic cells per mm²(median: 2,932, [range: 1,604-3,983]).

**Figure 5.**
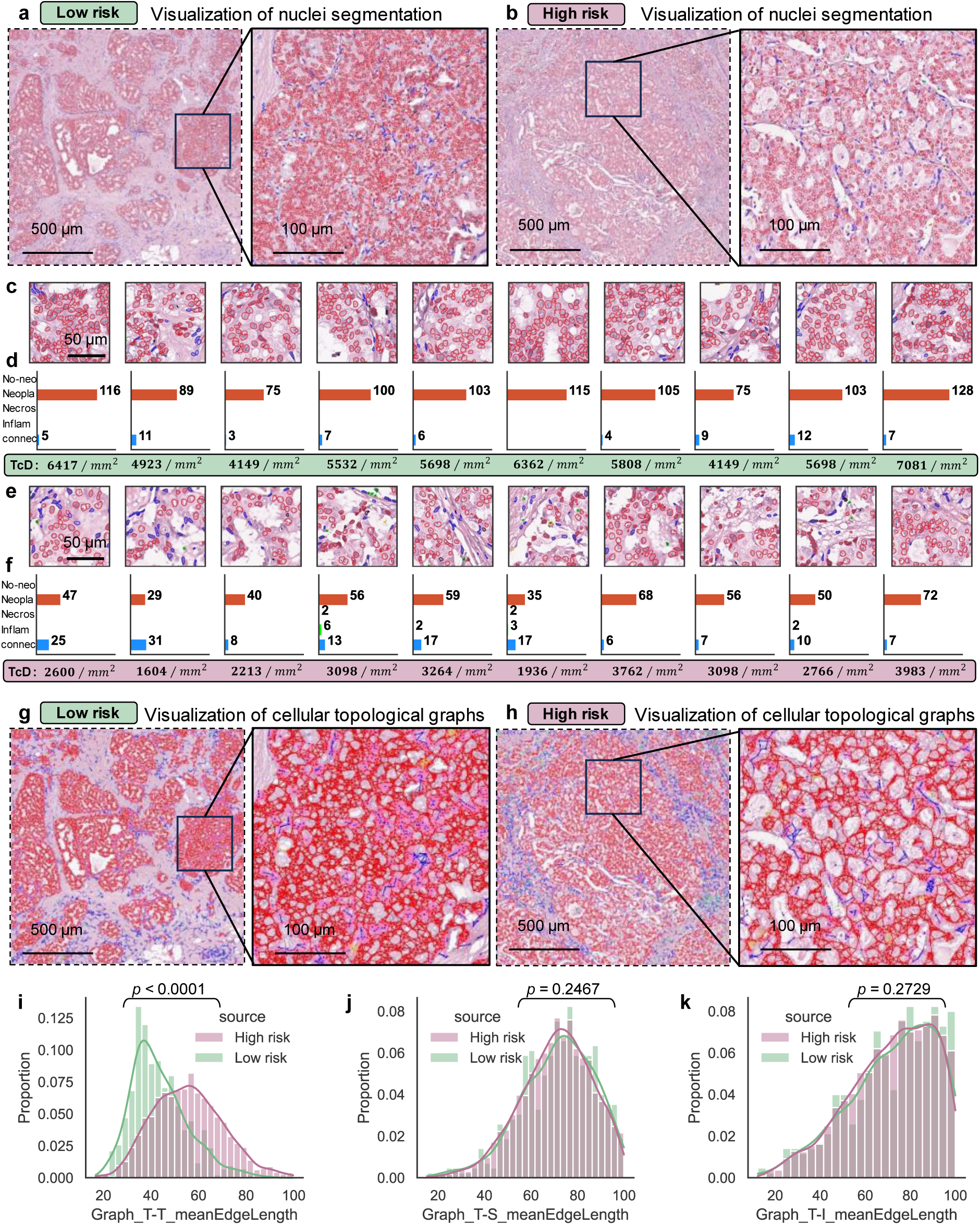
Intercellular interaction patterns within the tumor ecosystem. a, **b**. Visualization of nuclei segmentation results for MP Low-risk (**a**), High-risk (**b**) ROIs and their corresponding fine-grained regions. HoverNet was applied for simultaneous nuclei segmentation and classification. Each cell nucleus was classified as one of 5 cell types (neoplastic, inflammatory, necrotic, connective, and non-neoplastic epithelial cells). The contours of identified nuclei were located and drawn on the fine-grained pathological ROIs. Scale bar: 500 μm and 100 μm. **c, e.** Nuclei segmentation results of the top 10 local tiles with the highest attention scores for MP Low-risk (**c**) and High-risk (**e**). Scale bar: 50 μm. **d**, **f.** The cellular composition quantitation of the five cell types in each local tile region. The number of cells for each type is labelled on the right side of the plot, and the TcD values representing the density of tumor cells are marked in the lower part of the plot. **g, h**. Visualization of cellular interaction topological graphs in the tumor ecosystem. Three main cellular components (neoplastic, inflammatory, and connective stromal cells) within the tumor ecosystem were retained to construct topological graphs among these detected cell nuclei. MP Low-risk (**g**), High-risk (**h**). Scale bar: 500 μm and 100 μm. **i-k**. The feature distribution of graph-edge-length-related MeanEdgeLength (the average edge lengths of a tumor cell interacting with tumor cells (**i**, Graph_T-T_MeanEdgeLength), stromal cells (**j**, Graph_T- S_MeanEdgeLength), and inflammatory cells (**k**, Graph_T-I_MeanEdgeLength)) at the single-cell level for both MP risk groups. The sc-MTOP framework was leveraged for profiling the graph-based features of tumor cells interacting with other cells. *P*-value was calculated by the nonparametric Mann-Whitney U rank test. WSI=Whole Slide Image, ROI=Region of Interest, MP=MammaPrint diagnostic test, TcD=Tumor cell Density.

Further, neoplastic cells, connective cells, and inflammatory cells (the main cellular components in the tumor ecosystem^[46]^) were employed to construct topological graphs among these cell nuclei to facilitate our understanding of intercellular interactions within the tumor microenvironment (Figure 5g, h). The cellular topological graphs were visually characterized by interactions between tumor cells and stromal cells, with minimal evidence of lymphocyte presence and interaction with other cell types within both Low-risk and High-risk ROIs. To quantitatively characterize the spatial intercellular interaction patterns within the tumor ecosystem, the sc-MTOP^[47]^ framework was utilized to extract the topological graph-based features at the single-cell level. We only focus on profiling the topological features of tumor cells that interact with other cells, specifically examining the connectivity graphs of tumor-tumor, tumor-stroma, and tumor-inflammatory interactions (Figure S10b). The distribution of the MeanEdgeLength features (the average edge lengths of a cell interacting with other cells) related to graph-edge lengths of tumor cells was analyzed and depicted in Figure 5i-k. Among Low-risk patients, the graph-edge length distribution of tumor-tumor interaction graphs exhibits a higher peak, suggesting a denser and more abundant distribution of tumor cells. In contrast, High-risk patients display a sparse distribution of tumor cells (Figure 5i, *p* < 0.0001, statistically significant difference). The edge length distribution for tumor-stroma and tumor-inflammatory interaction graphs remains generally consistent across both patient groups (Figure 5j, *p* = 0.2467, and Figure 5k, *p* = 0.2729, no statistically significant difference). These results provide evidence that CPMP harnesses whole-slide-level heatmap visualization for deciphering the patterns of intercellular interactions within the tumor ecosystem, which are associated with genomic MP recurrence risk groups.

### 2.6. CPMP characterizes morphological phenotypes associated with MP risk groups

Breast cancer ecosystems often exhibit both intra-tumoral and inter-tumoral heterogeneity, which influences tumor progression, metastasis formation, and patient prognosis^[48]^. To this end, we aimed to characterize the tumor morphological patterns throughout the entire test set and explore the specific and common phenotypes associated with MP risk groups using CPMP. Specifically, the top 100 tiles with the highest attention scores in each slide were identified to create a collection of high-confidence local regions. Each tile in this collection was also fed into CPMP for inference. It can be observed from the uniform manifold approximation and projection (UMAP) plot that tile instances display a clear distinction among different levels of the genomic MP risk (Figure S11a). The recurrence risk prediction at both the WSI-level and the single-tile-level exhibits similar distribution characteristics in the UMAP (Figure S11b, c), in keeping with the observations from Figure S11a. The model’s capability of predicting recurrence risk at the single-tile level is more pronounced, as it demonstrates a clear trajectory direction of recurrence risk development (indicated by the arrow in Figure S11c). These results suggest that the high-confidence local tiles identified by the CPMP model provide a sufficient representation of WSIs.

We further performed Leiden clustering on tile embeddings to characterize phenotypic diversity of tumor morphology and attached the Low-risk or High-risk group information for each subcluster based on the genomic MP risk group of the WSI from which the local tile originated. High-attention tumor tiles were classified into 27 subclusters (named SC0-SC26) with different morphological embeddings (Figure S11d). Also, some subclusters are only present in a specific risk group, and others appear in both risk groups (**Figure 6**a-c and S12a, b). Our model identifies four subclusters (SC14, SC21, SC23, and SC24) that are exclusively associated with the MP High-risk group (Figure 6a) and 10 subclusters (SC1, SC5, SC6, SC7, SC9, SC10, SC20, SC22, SC25, and SC26) specifically related to the MP Low-risk group (Figure 6c). The remaining 13 subclusters colocalize in both MP risk groups (Figure 6b), reflecting the consistent tumor morphological characteristics of early-stage BC patients to whom the genomic MP test is applicable.

**Figure 6.**
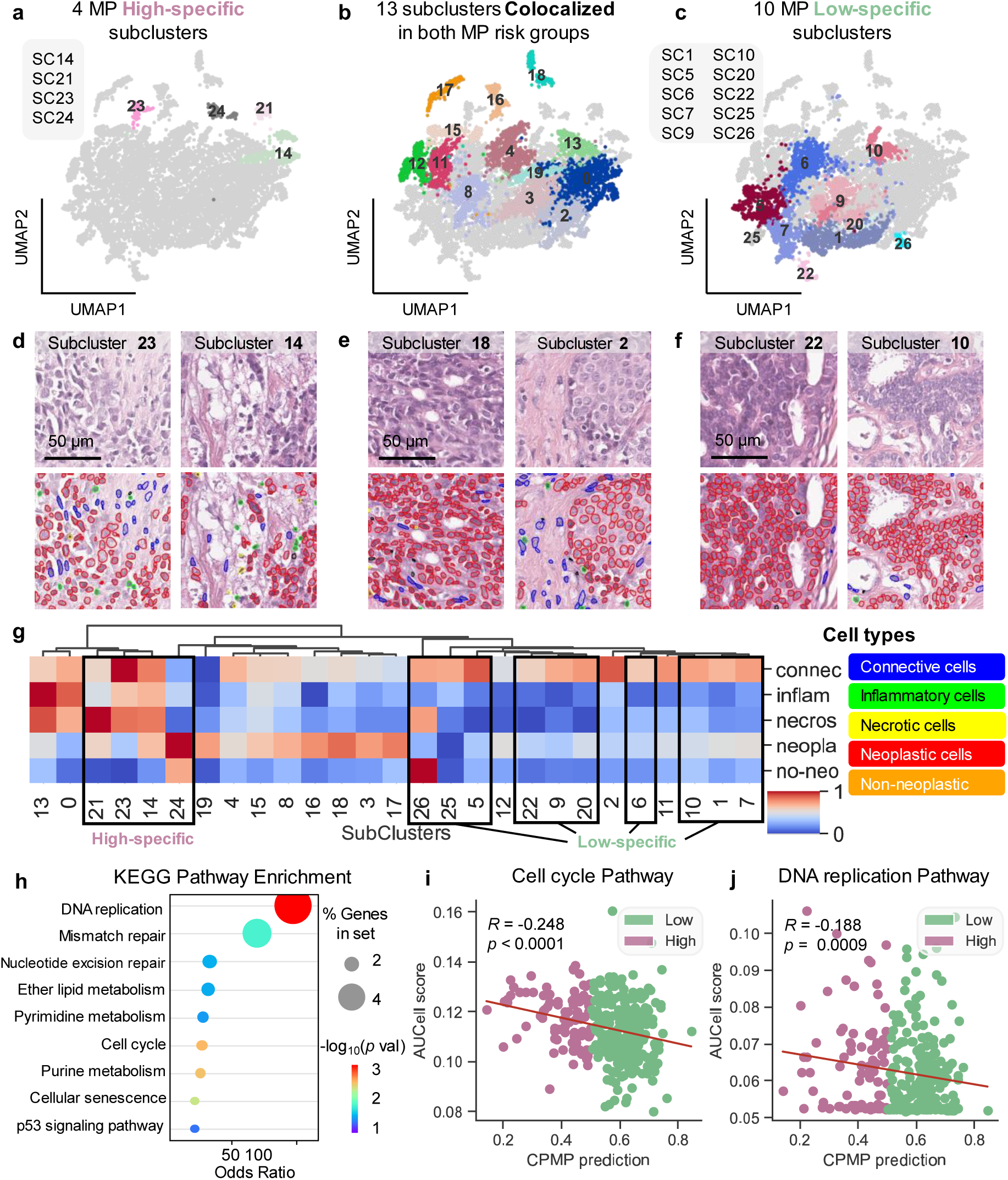
**Characterization of MP-specific and colocalized morphological phenotypes and their associated biological features. a-c**. UMAP subplots of High-specific (**a**), colocalized (**b**), and Low-specific (**c**) subclusters of embeddings of local tiles, derived from Leiden clustering of tile-level representations. Subclusters are labeled with their inferred IDs (SC0-SC26). The High-specific subclusters (**a**) are exclusively associated with the MP High-risk group, while the Low-specific subclusters (**c**) are specifically related to the MP Low-risk group. **d-f**. Depiction of representative H&E-stained local tiles and corresponding nuclei segmentation results for selected subclusters demonstrating the phenotypic characteristics of MP risk groups. (**d**) Subcluster 23 and 14 (High-specific), (**e**) Subcluster 18 and 2 (colocalized), and (**f**) Subcluster 22 and 10 (Low-specific). Nuclei are classified as connective (blue), inflammatory (green), necrotic (yellow), neoplastic (red), and non-neoplastic epithelial cells (orange). Scale bars: 50 μm. **g.** Heatmap depicting the cellular composition quantitation of the five cell types (connective, inflammatory, necrotic, neoplastic, and non-neoplastic epithelial cells) across the identified 27 subclusters. Min-max normalization was applied to scale the quantitative features of the cellular composition. Hierarchical clustering was performed on 27 subclusters using the Ward variance minimization algorithm and the Euclidean distance metric. Hierarchical clustering reveals distinct cellular compositions among subclusters, with High-specific and Low-specific subclusters forming separate clusters. **h**. KEGG pathway enrichment analysis of genes within the MP signature that showed a significant correlation with CPMP-predicted risk scores, including nine significantly enriched pathways. **i, j**. Scatter plots showing the correlation between CPMP-predicted risk score (x-axis) and AUCell scores (y-axis) for two key biological pathways: (**i**) Cell cycle and (**j**) DNA replication. The data points are colored by the corresponding genomic MP risk group (Low-risk in Spring green, High-risk in Lilac). The Spearman correlation coefficient (R) and *p*-value are reported for each pathway. The fitted linear regression line is also shown. A negative correlation is observed, suggesting that CPMP-predicted "High-risk" scores are associated with higher activity in these proliferation-related pathways. UMAP=Uniform Manifold Approximation and Projection, MP=MammaPrint diagnostic test.

Representative local tiles from each subcluster, along with the corresponding nuclear identification results, were depicted to highlight the morphological and phenotypic characteristics linked to genomic MP risk groups (Figure 6d-f, S12c, d, and S13a). The MP High-specific subcluster phenotypes (Figure 6d and S12c) exhibit loosely distributed tumor cells interconnected with stromal cells, while the MP Low-specific subcluster phenotypes (Figure 6f and S12d) present a high percentage of aggregated tumor cells. These visually observed morphological patterns are also reflected in the colocalized subcluster phenotypes (Figure 6e and S13a). These results reinforce that the morphological differences among subcluster phenotypes mainly depend on the patterns of intercellular interactions and the aggregation abundance of tumor cells. We next quantified the cellular composition of the five cell types for each subcluster and performed hierarchical clustering on the 27 subclusters (Figure 6g and S13b). Four MP High-specific subclusters are grouped together, while the MP Low-specific subclusters are clustered at the opposite end, forming four subgroups (SC26, SC25, and SC5; SC22, SC9, and SC20; SC6; and SC10, SC1, and SC7). The distinct cellular compositions between the High-specific and Low-specific subcluster phenotypes underline the capacity of CPMP to disclose the morphological patterns related to molecular risk groups.

To further explore the potential link between morphological features in the tumor ecosystem and underlying molecular biology mechanisms, we performed a KEGG pathway enrichment analysis on the subset of MP genes that showed a correlation with the CPMP- predicted risk score. Our analysis revealed that the most significantly enriched pathways (p- value < 0.05) include DNA replication, Cell cycle, Purine metabolism, Cellular senescence, Mismatch repair, Nucleotide excision repair, Ether lipid metabolism, Pyrimidine metabolism, and p53 signaling pathway (Figure 6h). These are consistent with the known biological functions^[49]^ (growth, proliferation, transformation, and cell death) of the 70-gene MP profile. We then analyzed the correlation between CPMP-predicted risk scores and pathway activity scores for each significantly enriched pathway (Figure 6i-j and S14). Significant negative correlations were observed for Cell cycle (R = -0.248, *p*-value < 0.0001) and DNA replication (R = -0.188, *p*-value = 0.0009), indicating that High-risk patients are characterized by heightened cellular proliferation at the molecular level. This micro-genomic proliferative activity presents co-occurrence with loosely distributed tumor macro-morphology at the whole-slide level among High-risk patients in our cohort.

### 2.7. CPMP serves as a potential tool for prognostic risk assessment

The predictive ability and generalizability of CPMP require validation in external cohorts. To this end, we collected and analyzed a new independent external cohort comprising 54 early-stage HR+/HER2− BC patients (20 High-risk, 34 Low-risk) from Tianjin Medical University Cancer Institute and Hospital, Binhai Hospital (TJMUCH-BH), which was processed using the same scanning platform as the TJMUCH-MP cohort. Without any model fine-tuning, we applied the trained CPMP model to this independent cohort for external evaluation. The model achieved an AUROC of 0.772 and showed a positive correlation (Spearman R = 0.433, p = 0.001) between genomic MP risk values and CPMP-predicted recurrence risk scores (Figure S15). These results provide quantitative evidence of CPMP’s robustness across clinical cohorts.

To further assess generalizability, we also evaluated CPMP in The Cancer Genome Atlas (TCGA)–BRCA cohort for recurrence risk and prognostic analysis (Figure S16a-c). This analysis was motivated by the established influence of MP-related gene activity on tumor progression and its association with patient outcomes. Whole-slide-level spatial heatmaps generated by CPMP in the TCGA-BRCA cohort reveal its ability to capture tumor localization patterns that may be clinically relevant (**Figure 7**a, b, S17, and S18). We further assessed the prognostic value of CPMP using clinical outcome endpoints. Disease-Free Interval (DFI),

**Figure 7.**
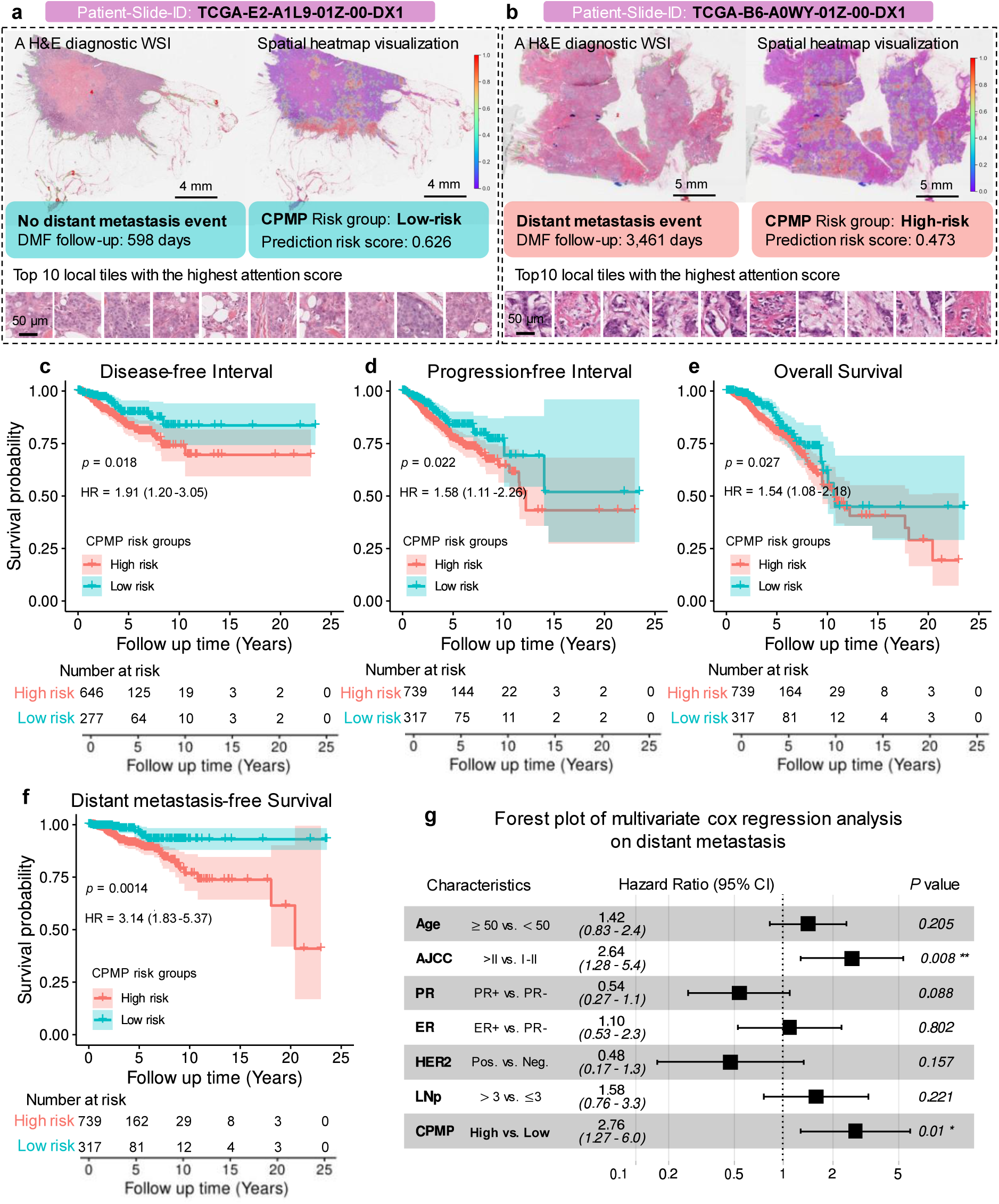
Prognostic risk stratification performance of CPMP on the external TCGA-BRCA data. a, **b.** Whole-slide-level spatial attention heatmap visualization results for tumor morphological patterns associated with prognostic events in the TCGA-BRCA dataset. The H&E-stained diagnostic WSIs of patients with (**b,** Scale bar: 5 mm) or without (**a,** Scale bar: 4 mm) distant metastasis event, the whole-slide-level overlaid spatial heatmap visualization derived from CPMP, and top 10 local tiles (Scale bar: 50 μm) with the highest attention score revealing the tumor morphological patterns of the slide are all shown. **c-f**. Kaplan–Meier curves for Disease-free Interval (**c**), Progression-free Interval (**d**), Overall Survival (**e**), and Distant metastasis-free Survival (**f**) of patient groups with High-risk and Low-risk in the external TCGA-BRCA dataset. Risk groups were directly predicted by CPMP trained in the TJMUCH-MP cohort. HR and 95% CI were calculated by the Cox proportional hazard model, while the *p*-value was calculated using the two-sided log-rank test. **g.** Forest plot of multivariate Cox regression analysis including the covariates age, AJCC pathologic tumor stage, PR status, ER status, HER2 status, LNp, and CPMP risk group, and their association with distant metastasis-free survival. HR and 95% CI were calculated by the Cox proportional hazard model, and the *p*-value was calculated using the two-sided log-rank test. (* indicates statistical significance, * p<0.05, ** p<0.01). WSI=Whole Slide Image, HR=Hazard Ratio, DMFS=Distant Metastasis-Free Survival, CI=Confidence Intervals, AJCC=The American Joint Committee on Cancer, PR=Progesterone Receptor, ER=Estrogen Receptor, HER2=Human Epidermal growth factor Receptor 2, LNp=The number of Lymph Nodes positive, HR+ =Hormone Receptor positive, LN0=Lymph Nodes negative, LN1=The number of Lymph Nodes positive≤3.

Progression-Free Interval (PFI), and Overall Survival (OS) were derived from the TCGA Clinical Data Resource^[50]^. We can observe a significant risk-group stratification on DFI, PFI, and OS prognostic indicators (log-rank test, *p*-value < 0.05) from Kaplan-Meier survival curves (Figure 7c-e). Patients in the High-risk group had a worse outcome than patients predicted as Low-risk by CPMP, exhibiting statistically significant findings in DFI (a hazard ratio (HR) of 1.91, 95% confidence interval (95% CI): 1.20-3.05, *p*-value = 0.018), PFI (HR: 1.58, 95% CI: 1.11-2.26, *p*-value = 0.022), and OS (HR: 1.54, 95% CI: 1.08-2.18, *p*-value = 0.027). We performed a prognostic analysis of CPMP risk groups correlating with Distant Metastasis-Free Survival (DMFS) in the TCGA-BRCA dataset (Figure 7f). Patients in the High-risk group experienced a higher probability of distant metastasis events than those classified as Low-risk (HR: 3.14, 95% CI: 1.83-5.37, *p*-value = 0.0014). Especially, the distinction between patients in the High-risk and Low-risk groups became more evident beyond the early survival times (∼ 5 years) as observed in the Kaplan-Meier (KM) survival curves for DMFS. This reveals the capability of CPMP in identifying late distant metastasis events. We investigated 16 patients suffering from distant metastasis beyond 5 years, of whom 14 (87.5%, a mean follow-up time of ∼9.44 years) were identified as High-risk by CPMP. We also evaluated the prognostic impact of the predicted risk groups on DMFS, DFI, PFI, and OS using multivariable Cox proportional hazards models (Figure 7g and S19a-c). The risk groups inferred by CPMP exhibited a significant prognostic impact on DMFS, with an HR of 2.76 (95% CI: 1.27-6.00, *p*-value = 0.01, Figure 7g). It suggests that the CPMP-derived risk group is an increased risk factor for BC distant metastasis in TCGA-BRCA patients, demonstrating the prognostic capability of CPMP. It can be implied that CPMP effectively captures valuable prognostic features from tumor morphology in BC histopathological slides. This further highlights the potential of CPMP to be applied in clinical settings for prognostic risk assessment of BC patients.

## 3. Discussion

In this study, we presented the weakly supervised CPMP model for MP recurrence risk prediction from H&E-stained histopathological WSIs of BC patients. The predictive capability of the CPMP model was shown by inferring both the continuous risk scores and the discrete recurrence risk groups in the real-world TJMUCH-MP clinical cohort. Compared to the existing weakly supervised MIL methods, our method achieved superior AUC-efficiency performance and yielded a better alignment with genomic MP risk scores in the cohort.

To the best of our knowledge, this is the first work to explore spatially morphological tumor patterns associated with genomic MP for recurrence risk assessment and prognosis analysis in BC patients. Our model facilitated the characterization of the tumor pattern diversity in early-stage BC patients for whom the genomic MP test is applicable. It enhanced the discovery of High-specific, Low-specific, and colocalized morphological phenotypes among MP risk groups from routine histopathology slides. The intercellular interactions and quantized cellular composition patterns exhibited distinctions across diverse tumor phenotypes, which are related to genomic MP recurrence risk groups. Moreover, our study provided evidence that the CPMP model demonstrated superior generalization capabilities in identifying late distant metastasis events in the TCGA-BRCA cohort. It also exhibited significant risk-group stratification capabilities across multiple prognostic indicators. We propose that the prognostic value of CPMP arises from its ability to bridge the gap between macro-morphology and micro-genomic expression, by capturing the potential connections between spatial tumor morphology and underlying gene activities.

The genomic MP test provides significant clinical value and has been proven to be an effective diagnostic tool in current clinical applications. It is recommended for invasive BC patients who are HR+/HER2-, staged T1-2, and either lymph node-negative or with 1-3 metastases^[29]^. This test evaluates the 10-year risk of distant metastasis and classifies patients into High or Low recurrence risk groups. High-risk patients are likely to benefit from adjuvant chemotherapy, while Low-risk patients with clinically high-risk features can safely be exempted from chemotherapy. The genomic MP profiling test for recurrence risk diagnosis is performed on tumor specimens using targeted RNA next-generation sequencing technology. However, it fails to provide the spatial and morphological characteristics of tumors. To make up for the limitations of traditional MP diagnostic tests in terms of spatial analysis, we implemented a WSI-level attention heatmap visualization method in our model. This approach captures the spatial relationships and cellular interaction patterns within tumors at the whole-slide level. This indicates that our spatially aware, weakly supervised learning CPMP model is capable of generating a whole-slide-level attention heatmap, facilitating the visualization and discovery of spatial morphological localization.

In addition, the time and financial costs of MP diagnostic tests are substantial, creating a barrier that prevents many patients from accessing precise therapeutic regimens. CPMP, an AI-driven weakly supervised learning model, demonstrates remarkable capability in predicting distant metastasis risk for early-stage BC patients using only routine histopathological slides. We experimentally validated that the CPMP model can infer a reliable risk score and a whole-slide-level spatial attention heatmap visualization within approximately 5 minutes for a WSI of size 50,000×50,000 pixels (25 mm × 25 mm). This approach not only reduces the financial burden on patients but also enables rapid result delivery. Consequently, CPMP presents a viable complementary option for patients who cannot afford the genomic MP test and has the potential to serve as a cost-efficient, AI-powered system for pre-screening BC patients in the clinical workflow (depicted in Figure S20).

Our study brings favorable insights into the fields of computational pathology and biomarker discovery, but there are still certain limitations. For example, our weakly supervised CPMP model leverages general-purpose foundation models^[39–44]^ for feature extraction, which reduces computational costs. However, these foundation models were pretrained on large datasets whose data distribution may differ from ours due to variations in slides. As a result, one line of future work is to address these domain distribution challenges by fine-tuning existing self-supervised models^[51,52]^. While this approach has the potential to enhance predictive performance, the associated computational resources must be considered, and the trade-off between performance gain and resource cost needs careful evaluation. Additionally, our model was developed using the TJMUCH-MP cohort and is currently limited to predicting MP High and Low risk groups for early-stage BC patients. Recent studies^[53,54]^ suggest that an Ultra-High (H2) subgroup exists within the High-risk group, potentially benefiting from immunotherapy. However, the number of H2 patients in the TJMUCH-MP cohort is limited (n = 11), which constrains the CPMP’s ability to learn and identify this subgroup. We will expand the cohort through multi-institutional collaborations to develop and validate a refined model capable of identifying this high-impact patient population. Furthermore, although we have demonstrated the model’s generalization capability in the external TCGA-BRCA cohort, its robustness to domain shifts caused by variations in slide preparation, staining, and scanning protocols across different institutions and ethnic populations remains incompletely validated. For broader clinical application, multi-center clinical trial studies involving diverse, multi-ethnic cohorts will be essential to evaluate CPMP’s real-world impact on clinical outcomes such as treatment decisions and patient survival. To improve cross-institutional robustness, future efforts could incorporate methodological advancements in domain adaptation. For instance, unsupervised domain adaptation^[55]^ can be leveraged to align the feature distributions of histopathological images between the source domain and new target domains, thereby mitigating domain shifts. Alternatively, self-supervised fine-tuning on a small set of unlabeled WSIs from a target institution may help the model adapt to local staining and scanning protocols. In parallel, future research should aim to further elucidate the intercellular morphological patterns associated with MP recurrence risk groups and investigate the causal links between tumor morphology and the MP genomic signature.

Altogether, our study revealed that AI-driven weakly supervised computational pathology technology can effectively predict risk scores for prognostic distant metastasis hints using annotation-free histopathological WSIs. This capability can further facilitate prognostic knowledge discovery, paving the way for advancements in precision treatment. These findings make it possible to achieve a complementary, computational, and cost-efficient diagnostic tool for assessing recurrence risk for early-stage BC patients from routine H&E-stained slides, ultimately supporting better clinical decision-making and patient management.

## 4. Methods

### 4.1. Clinical cohort curation

The TJMUCH-MP cohort examined female patients who were diagnosed with primary breast cancer and received treatment between January 2019 and December 2023 at TJMUCH, National Clinical Research Center for Cancer, China. Patients who underwent MP diagnostic tests were included in this retrospective analysis. Whole-slide images were scanned using the Aperio Leica Biosystems GT450 v1.0.1 scanner at an apparent magnification of 40×. The genomic MP test was conducted by ZhenHe Genecast Biotechnology Ltd, the exclusive partner for the 70-GS assay in China, as appointed by Agendia. All participants enrolled in this study met the StGallen guidelines for the MP test^[29]^. Inclusion criteria comprised invasive BC that was HR+/HER2-, staged T1-2, and either lymph node-negative or with 1-3 metastases. Additionally, tumor specimens were required for genomic analysis. Exclusion criteria included HR- or HER2+ tumors and patients receiving neoadjuvant therapy. Cases involving bilateral breast cancer and inconclusive tissue sections were also excluded. Clinical information and histopathological characteristics were extracted from medical records, while adverse events, such as recurrence and death, were tracked through follow-up visits. FFPE tumor samples were collected from biopsies and excised surgical materials. Based on the genomic MP diagnostic test, continuous risk scores were yielded, ranging from -1 to 1. Patients were categorized into High-risk and Low-risk groups using a cutoff value of 0.0, with scores ≤ 0.0 categorized as High-risk and scores > 0.0 as Low-risk^[7]^. For model training and prediction, the original MP risk scores were normalized to the range [0, 1] using the formula (x + 1) / 2. After normalization, a threshold of 0.5 was applied for binary classification, where scores ≤ 0.5 were classified as High-risk and scores > 0.5 as Low-risk. Ultimately, a HR+/HER2- invasive BC cohort was established, comprising digital WSIs, genomic MP results, and clinicopathological features of 477 female patients (Figure S1a-d, S2a-b, and Table S1). This cohort was utilized for subsequent model development and evaluation.

### 4.2. WSIs preprocessing

Histopathological WSIs typically contain billions of pixels and possess high-resolution, multi- scale attributes. However, these images include areas of white background and non-tissue regions that should be filtered out. To improve the computational efficiency of WSI analysis, we first performed data preprocessing on the H&E-stained histopathological slides (Figure S2c). We utilized automated tissue segmentation techniques proposed by CLAM^[35]^ for effective tissue detection, resulting in segmented foreground contours. Following this, we employed a non-overlapping sliding window tiling strategy to crop image tiles measuring 256×256 pixels from the identified tissue regions at a 20×magnification. The resulting stacks of image tiles, along with their spatial coordinates, were organized and stored using the Hierarchical Data Format version 5 (HDF5)^[35]^. The distribution of the number of tiles obtained from each WSI varies widely, ranging from thousands to hundreds of thousands, as illustrated in Figure S2d.

### 4.3. CPMP model

Our model employs the weakly supervised MIL framework for predicting recurrence risk from histopathological WSIs (Figure 1a). To clarify, we begin with outlining the classic MIL approach. In this framework, a dataset 𝔻 = {(𝒳_𝒾_ 𝒴_𝒾_)}^𝑁^ consists of a total of 𝑁 bags 𝒳_𝒾_, each containing multiple instances 𝒳 = {𝑡^𝒾^}^𝓃𝒾^ . The primary objective of the MIL approach is to develop a function ℱ(·) that can predict the bag-level category 𝒴_𝒾_ based on the multiple instances {𝑡^𝒾^}^𝓃𝒾^ within that bag. While the bag-level labels {𝒴 } are known, the labels for the individual instances {𝑡^𝒾^} are unknown. The function is typically decomposed into three key components: 1) An instance-level transformation function that transforms instances into feature vectors. 2) An aggregation function at the instance-level that integrates the features of all instances to yield a global feature representation at the bag-level. 3) A classifier head function that predicts the bag-level results from this global feature representation. In alignment with the MIL paradigm, CPMP is composed of three components: a feature representation function 𝒯(·) operating at the tile-level, a feature aggregation function 𝒮(·) that integrates features across tiles within a slide, and a regression-based MLP predictor head ℛ(·) at the slide-level. We describe these components in detail in the following section.

#### 4.3.1. Feature representation at the tile-level

Feature representation, serving as the instance-level transformation function in MIL, was executed on tiles cropped from WSIs. Recently, several self-supervised models^[39–44]^ pretrained on extensive datasets of histopathological WSIs across pan-cancer pan-tissue types have become publicly available, significantly facilitating the advances in the field of computational pathology. These general-purpose foundation models succeed in capturing the diverse patterns in pathology images and hold promise for integration into clinical diagnostical workflows, particularly in applications with limited data availability. We explored several models, including UNI^[41]^, Prov-GigaPath^[39]^, Virchow^[40]^, H-optimus-0^[42]^, Phikon^[44]^, and CTransPath^[43]^, for tile-level feature extraction (Figure S2e). We first resized the 256×256 tiles to 224×224 pixels before inputting them into feature encoders for forward computation. This process allows us to represent each tile with low-dimensional feature vectors: 1024-dimensional (UNI encoder), 1536-dimensional (Prov-GigaPath), 2560- dimensional (Virchow), 1536-dimensional (H-optimus-0), and 768-dimensional (Phikon and CTransPath). Low-dimensional feature representations at the tile level make it feasible to reduce computational complexity and aggregate diverse patterns from all tiles within a slide.

#### 4.3.2. Feature aggregation

Based on the feature embeddings of local tiles, we employed the Transformer^[56]^ architecture for feature aggregation to derive a global feature representation at the slide-level. The self-attention module in the Transformer is crucial, as it can capture long-range dependencies across tiles. However, the global self-attention mechanism incurs a high computational cost, particularly when the number of tiles in a single slide can reach up to 100,000. This excessive computation and redundancy limit the applicability of the self-attention mechanism in our high-resolution scenarios. To this end, we implemented an agent-attention mechanism^[30]^ to integrate the embedding sequences by introducing an agent token (Figure 1b, c). This modification transforms the traditional self-attention (𝒬 𝒦 𝒱) into the agent-attention paradigm (𝒜 𝒬 𝒦 𝒱), balancing between the need for global representation capabilities and the requirement for computational efficiency.

Specifically, for the input tile features 𝑥 = {𝒯(𝑡)}^𝓃^ 𝑥 ∈ ℝ^𝓃×𝑐^, the general self-attention module can be formulated as follows:

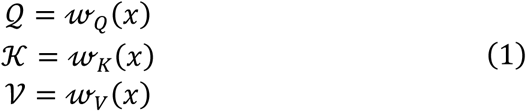

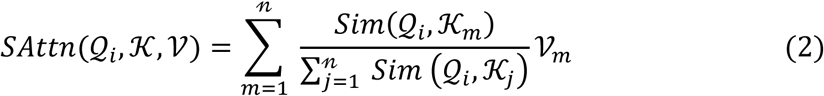

where 𝓃 and 𝑐 are the number of tile embeddings in one slide and the dimension of tile embedding, respectively. 𝓌_𝑄_ 𝓌_𝐾_ 𝓌_𝑉_ ∈ ℝ^𝑐×𝑑^denote respective linear transformation layers for Query, Key, and Value in the Transformer block, where 𝑑 is the output dimension of linear layers. 𝑆𝑖𝑚(·) represents the similarity function, commonly 𝑆𝑖𝑚(𝒬 the Softmax attention. We abbreviate the similarity function to 𝜎(·) and the Softmax attention can be rephrased as follows:

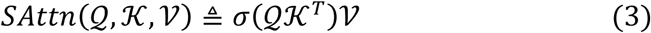

Softmax attention computes a query-key product, resulting in a computation complexity of 𝒪(𝑛^2^). It is noteworthy that we do not consider multi-heads here.

In the agent-attention paradigm, we added an agent token 𝒜 to the foregoing attention triplet (𝒬 𝒦 𝒱). This modification results in the agent-attention paradigm of (𝒜 𝒬 𝒦 𝒱), which we denote as 𝐴𝐴𝑡𝑡𝑛(𝒜 𝒬 𝒦 𝒱). To be specific, the agent-attention paradigm consists of a stacked state of two Softmax attention operations 𝑆𝐴𝑡𝑡𝑛(·). In the first phase, referred to as 1st-aggregation, the agent token 𝒜 is treated as a Query 𝒬 in the first Softmax attention operation 𝑆𝐴𝑡𝑡𝑛(𝒜 𝒦 𝒱). Information is aggregated into the Agent-Value 𝒱_𝑎𝑔𝑒𝑛𝑡_ from the Value 𝒱, with the attention matrix computed based on Agent 𝒜 and Key 𝒦. The second Softmax attention operation 𝑆𝐴𝑡𝑡𝑛(𝒬 𝒜 𝒱_𝑎𝑔𝑒𝑛𝑡_) is applied in the second phase, known as 2nd-broadcast. Here, the Agent-Value 𝒱_𝑎𝑔𝑒𝑛𝑡_ serves as the output from the first Softmax attention. This process yields the final output from the agent-attention mechanism (Figure 1c).

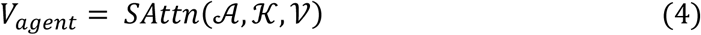

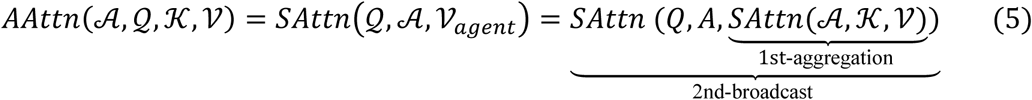

The pairwise Query-Key product computation is replaced by the Agent-Key and Query- Agent product, and the information flow is connected and preserved through the Agent. Notably, the number of agent tokens 𝑘 is a very small constant, which theoretically reduces the computational complexity of the agent-attention to 𝒪(𝑛𝑑), where 𝑑 ≪ 𝑛.

Using the agent attention mechanism described above, we employed the agent-attention transformer for instance aggregation (Figure 1b). We first transformed 𝓃 tile embeddings into a dimension of 256 through a linear projection layer, followed by the non-linear activation function ReLU. To incorporate the spatial positional information of the tiles, we employed a 2-dimensional absolute position encoding method^[56,57]^ to encode the tile coordinates. The resulting 256-dimensional positional embeddings were then added to the 256-dimensional tile embeddings. Next, a class token 𝑥_𝑐𝑙𝑠_ ∈ ℝ^1×256^was initialized and concatenated with the spatially-aware tile embeddings, yielding a fused representation {𝑥 𝑥_𝑐𝑙𝑠_} ∈ ℝ^(𝓃+1)×256^. We configured an agent-attention transformer block consisting of one layer with four heads for global information perception and aggregation. The output dimensions of the latent projection layers for Query, Key, and Value were set to 256, while the Feed Forward hidden linear projection layers produced an output dimension of 512.

#### 4.3.3. A regression-based MLP predictor head

Based on the agent-attention transformer block, the class token 𝑥_𝑐𝑙𝑠_was utilized as the aggregated global representation at the slide-level. This token was then passed into the MLP predictor head, which included a LayerNorm normalization followed by a linear layer. Given that the patient’s MP risk is represented as a continuous score, binarizing these values may result in information loss. Recent studies^[58,59]^ provide evidence that regression-based approaches are superior to classification-based methods for predicting continuous biomarkers. Therefore, we adopted a regression-based strategy to output a continuous numerical value as the risk prediction result. The linear projection layer transformed the feature embedding of the class token 𝑥_𝑐𝑙𝑠_into a single-neuron logit output, which was then passed through a Sigmoid activation function to yield a continuous risk value between 0 and 1. We employed the mean square error loss as the overall objective function ℒ(·) to optimize CPMP.

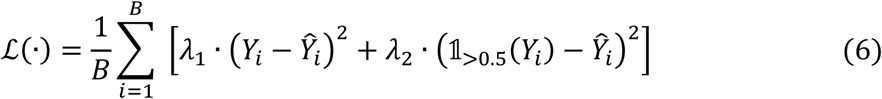

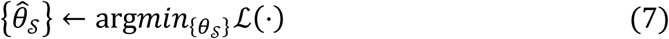

where 𝐵 is the batch size during training, 𝑌_𝑖_represents the slide-level continuous MP risk score, and 𝑌^_𝑖_ signifies the predicted continuous risk probability value for a patient. The parameters 𝜆_1_ and 𝜆_2_ are scale factors that balance the contributions of different loss components in the objective function. The indicator function 𝟙_>0.5_(·) returns 1 when 𝑌_𝑖_ ., and 0 when 𝑌_𝑖_ ≤ . . The optimization process is represented as arg𝑚𝑖𝑛_{𝜃𝒮}_ℒ(⋅), indicating that the optimization algorithm seeks to find the optimized parameters {𝜃_𝒮_} which minimize the objective loss ℒ(·). Here, {𝜃_𝒮_} refers to the set of optimized parameters in our model.

### 4.4. Spatial attention heatmap visualization at the whole-slide level

The whole-slide-level attention heatmap visualization was conducted to explore spatial localization in tumors. It was implemented by mapping attention scores of tiles to their corresponding locations within WSIs. Specifically, we first performed data preprocessing on each WSI to detect tissue areas. The identified foreground tissue regions were tessellated into 256×256 tiles with an overlap rate of 0.1 at a 20× objective magnification. Subsequently, we leveraged the self-supervised foundation model UNI for feature extraction from these tiles, resulting in 1024-dimensional feature vectors. The extracted tile embeddings were then passed through CPMP for inference, producing slide-level prediction results alongside tile-level attention scores that contributed to the final risk prediction. The attention scores at the tile-level were calculated using the attention rollout method^[60]^, based on our agent-attention matrix 𝑀_𝐴𝐴𝑡𝑡𝑛_(·) between tiles.

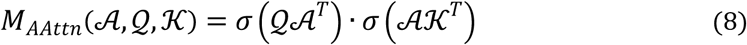

These attention scores were normalized to a range between 0 and 1, serving as attention weights. Finally, the normalized attention scores were arranged according to their corresponding spatial coordinates within the WSI, allowing for the visualization of an attention heatmap. This heatmap can be optionally overlaid on the original H&E-stained WSI, providing insights into the spatial distribution of attention that aligns with tumor areas.

### 4.5. Cell identification and intercellular graph construction

To gain further insights into the cellular interactions within the tumor ecosystem of MP High and Low risk groups, we first extracted the high-attention ROIs highlighted by the spatial attention heatmap at the whole-slide level. Subsequently, we applied HoverNet^[45]^ for simultaneous segmentation and classification of nuclei. We utilized the pretrained model weights trained on the public PanNuke dataset. The identified cells were classified into one of five cell categories: neoplastic (tumor), inflammatory, necrotic (dead), connective (stroma), and non-neoplastic epithelial cells. The spatial coordinates of the centroids and contours of these cells were located in the ROIs (illustrated in Figure S10a).

We defined the density of tumor cells (TcD) as the number of identified neoplastic cells per square millimeter. To further analyze the tumor ecosystem, we selected the three main cellular components^[46]^ (tumor, inflammatory, and stromal cells) to construct topological graphs among the detected cell nuclei. The sc-MTOP framework^[47]^ was employed for profiling graph-based features at the single-cell level within the tumor microenvironment. We focused on the topological features of tumor cells interacting with other cells, specifically examined the connectivity graphs of tumor-tumor, tumor-inflammatory and tumor-stroma interactions. Topological features related to graph-edge length, especially MeanEdgeLength^[47]^, were calculated to characterize the spatial intercellular relationships among different cell types within tumors. This metric represents the average edge lengths of each tumor cell interacting with other tumor, stromal, and inflammatory cells (illustrated in Figure S10b).

### 4.6. Characterization of phenotypic diversity

To investigate phenotypes potentially associated with MP recurrence risk groups, we first chose the model demonstrating optimal performance from the experiments based on a patient-level 5-Time 5-Fold cross-validation scheme. The histopathological WSIs of the testing cases were then tessellated into 256×256 tiles, which were characterized using the UNI foundation model for feature representation. These tiles were subsequently fed into the CPMP model for inference, yielding WSI-level risk scores. In addition, we identified and saved the top 100 tiles with the highest attention scores from the slide, which were deemed representative of local regions within the tumor. Each instance in this tile collection was also processed through CPMP to infer tile-level risk scores. The intermediate vector (cls_token) between the transformer block and the MLP classifier was detached, resulting in a 256-dimensional embedding vector for each tile instance.

The Scanpy toolkit was employed for further analysis of the embedding data. All tile embeddings were combined into an AnnData object, which contained a data matrix of the tile embeddings along with the additional metadata. The metadata included the spatial coordinates of the tiles within WSIs, genomic MP risk results corresponding to the slide from which the tile originated, WSI-level risk scores, and tile-level risk scores from the CPMP model. The nearest neighbors distance matrix and a neighborhood graph of tile embeddings were computed using the scanpy.pp.neighbors() function, employing cosine similarity as the distance measurement metric. The minimum size for local neighborhoods was set to 50 to balance between local information and global views. Based on the constructed neighborhood graph, Leiden clustering implemented by the scanpy.tl.leiden() function was performed for subcluster community detection on tile embeddings. Finally, we applied UMAP dimensionality reduction to visualize the tile embeddings and their subclusters in two-dimensional space via the scanpy.tl.umap() function. The low dimensional embedding was initialized using coarse-grained connectivity structures generated by the scanpy.tl.paga() function, which employs the partition-based graph abstraction (PAGA) algorithm.

Based on the Leiden clustering on the tile embeddings, we identified subclusters representing local morphological patterns. The following analytic steps were undertaken for these subclusters. First, we extracted the respective submaps for Low-risk and High-risk groups from the global subclusters, based on the MP risk group of the WSI from which the tile originated. We only preserved those subclusters whose number of tiles exceeds 25% of that in the corresponding global subcluster in a specific risk group. Next, we determined the intersection of subcluster ID sets from both the High- and Low-risk groups to identify subclusters that present in both MP risk categories, termed as colocalized subclusters. Subcluster IDs exclusive to the High-risk group were identified as High-specific subclusters, while those unique to the Low-risk group were designated as Low-specific subclusters. Subsequently, we applied HoverNet to all tiles within the High-specific, Low-specific, and colocalized subclusters for nuclei segmentation and classification. This process identified five cell types (neoplastic, inflammatory, necrotic, connective, and non-neoplastic epithelial cells), as well as their corresponding spatial coordinates within the tiles. Finally, we quantified the cellular composition for each subcluster by calculating the average number of cells for each cell type across all tiles within the subcluster. We then applied Min-max normalization to standardize these quantitative features of the cellular composition and conducted hierarchical clustering on all subclusters using the Ward variance minimization algorithm and the Euclidean distance metric.

### 4.7. Prognostic analysis on TCGA-BRCA

We applied the CPMP model to the external TCGA-BRCA cohort for recurrence risk assessment and prognostic analysis. TCGA-BRCA cohort includes 1,056 patients with digital diagnostic slides as well as clinical follow-up information (Figure S16a). All digital slides (n = 1,133) in the TCGA-BRCA cohort were first preprocessed and tessellated into local tiles. Tile-level feature representations were generated using UNI, and they were fed into the trained CPMP model for slide-level distant metastasis risk score inference. The prediction results of multiple slides from the same patient were averaged into the patient-level risk values. The continuous prediction risk scores were categorized into High-risk and Low-risk groups based on a cutoff value determined by the trained model. Spatial attention heatmap inference at the whole-slide level was also performed to visualize how regions with distinct tumor morphological patterns contribute to patient clinical outcomes. The prognostic value of the CPMP model within the TCGA-BRCA cohort was assessed using the Kaplan-Meier survival curves and the multivariable Cox proportional hazards model. The clinical outcome endpoints including Disease-Free Interval, Progression-Free Interval, and Overall Survival were derived from the TCGA Clinical Data Resource^[50]^.

### 4.8. Comparative methods

We compared the CPMP model with existing state-of-the-art weakly supervised MIL methods in WSIs classification. Brief introductions of the comparative methods are summarized as follows:

1. ABMIL^[34]^, an attention-based MIL method, currently a MIL baseline for WSI classification in the field of CPATH.
2. CLAM^[35]^, an instance-level clustering-constrained-attention MIL algorithm for pathological subtyping. It relied on instance-level clustering to refine the feature space in MIL.
3. DTFD-MIL^[33]^, a double-tier MIL framework that introduces pseudo-bags to enhance computational efficiency.
4. SAM-MIL^[32]^, a MIL framework that leverages the Segment Anything Model (SAM) to extract and integrate spatial context from WSIs into model training.
5. CEMIL^[31]^, an instructor-learner framework utilizing knowledge distillation for resource-efficient, low-computational-cost learning.
6. The full Transformer-based architecture model applied for end-to-end biomarker prediction from WSIs^[36]^ (denoted as Wagner et al.). We halved the architecture to a smaller scale with 4 attention heads due to data limitations in our experiments.
7. TransMIL^[37]^, a correlated Transformer-based MIL framework for pathology diagnosis on WSIs. It designed a PPEG for position encoding to capture both the local contextual information and the correlation between regions.

To verify the effectiveness of CPMP in using annotation-free WSIs, we conducted the experiments on noisy tissue filtering, tile sampling, and clinical-features-only logistic regression. In the experiment of tissue filtering, we aimed to investigate CPMP’s sensitivity to noisy tiles. We performed zero-shot classification to determine tissue types for all tiles using the visual-language foundation model PLIP^[38]^ (illustrated in Figure S5a). Tiles were identified as one of eight tissue types: tumor, adipose, stroma, immune infiltrates, gland, necrosis or hemorrhage, background or black, and non (Figure S5b). Tiles classified as “background or black” and “non-tissue” types were filtered out. In contrast, tiles identified as tumor, adipose, stroma, immune infiltrates, gland, and necrosis or hemorrhage tissue types were preserved and utilized for training the other model. This model with noisy tile filtering was compared with the CPMP model. To evaluate the scale flexibility of CPMP on the number of local tiles in WSIs, we conducted random sampling from the set of local tiles in each WSI, employing sample sizes of 1,000, 5,000, and 10,000 tiles as subsets for model training. The trained models were further evaluated on multiple subsets of tiles with variant sizes.

A clinical-based logistic regression model designed to predict patients’ MP recurrence risk from clinicopathological features was used for comparison. This clinical-based model was constructed using a Logistic Regression classifier, incorporating a set of eight clinical variables: ’Surgery Option’, ’T’, ’N’, ’AJCC’, ’Histological grading’, ’TILs’, ’PR’, and ’Ki67’. The clinical-based model underwent training, validation, and independent testing on the same patient cases, utilizing a patient-level 5-Time 5-Fold cross-validation strategy, consistent with the approach used in the CPMP model. Detailed hyperparameter settings for the Logistic Regression classifier are provided in the Supporting Information. Among the eight clinical variables, ’T’, ’N’, ’AJCC’, and ’Histological grading’ are categorical variables while ’TILs’, ’PR’, and ’Ki67’ are continuous variables representing intensity percentages or positive proportions, with values ranging from 0 to 100. The ’Surgery Option’ variable was categorized by assigning numerical values to different surgical options, with the specific categorization rules outlined in Table S5.

We also performed ablation studies to analyze modules in our spatially-aware, regression-based weakly supervised transformer architecture. The design outline and the experimental configurations utilized for the ablation models were illustrated in Figure S6a. Specifically, we first analyzed the transformer-based aggregation function with self-attention or agent-attention mechanisms. Regression-based mean square error (MSE) loss strategy was compared with classification-based cross-entropy (CE) loss. Positional coordinate information embedding (PE) was introduced into tile embeddings for spatial awareness in the CPMP model. PE, No-PE, and PPEG-based experimental configurations were thus adopted for comparison. In the ablation study of tile-level feature representation, we explored six self-supervised general-purpose foundation models, UNI^[41]^, Prov-GigaPath^[39]^, Virchow^[40]^, H-optimus-0^[42]^, Phikon^[44]^, and CTransPath^[43]^, for tile-level feature extraction and experimental comparison.

### 4.9. Implementation details

CPMP was implemented using Python 3.8 and the PyTorch 2.2 deep learning framework. All models, including comparative methods, were conducted on one workstation equipped with two 24GB NVIDIA GPUs (GeForce RTX 4090). The CUDA version is 12.2 and the GPU driver version is 535.171. Adam algorithm was employed for parameters optimization. The momentum factor was 0.9, and the learning rate was initially set to 0.0001. LinearLR scheduler aimed at enabling the learning rate decay was operated and set to a limit of 0.00001 when the epoch count reaches 500. The maximum number of epochs was set to 1000, and an early stopping strategy was utilized. The early stop is determined when the loss value of the validation set no longer decreases for 50 consecutive epochs. A gradient accumulation strategy was also adopted to address the issue of an inconsistent number of patches across WSIs. The gradient accumulation size was set to 32, which is equivalent to using a minibatch size of 32. The weights of all layers were initialized with the Kaiming uniform strategy. The optimized model with the lowest loss metric value on the validation set was recorded and utilized for evaluation on the test set. WSIs were processed by OpenSlide Toolkit v3.4.1.

For fair evaluation in case of data bias, we employed a patient-level 5-Time 5-Fold cross-validation scheme (Figure S3a) in the TJMUCH-MP cohort. In the experiment of each Time, we randomly divided 20% of the samples for testing, while the remaining samples were evenly divided into 5 non-overlapping subsets (Folds). Slides from each patient were only assigned to one of these sets to prevent data leakage. We then iteratively trained the model on 4 subsets (4 Folds) while using the remaining subset for validation. This process was repeated for each Fold, ensuring that each Fold served as the validation subset exactly once. The trained models corresponding to the 5 Folds were evaluated on the independent testing cases, and the results were averaged. This experiment was repeated for 5 random Times and the average results over the 5-Time experiments are reported.

### 4.10. Statistical analysis

The original genomic MP risk scores (ranging from -1 to 1) were normalized to the range [0, 1] using the formula (x + 1) / 2 for model training and prediction. This transformation was necessary to ensure compatibility. No other data transformation or normalization was applied. Data and results are presented as mean ± standard deviation (SD) where applicable, or as median and range for non-normally distributed data (e.g., tumor cell density). The sample size for the performance evaluation was n = 477 (TJMUCH-MP cohort). For survival analysis in the TCGA-BRCA cohort, the sample size was n = 1,056 patients. The nonparametric Spearman rank-order correlation coefficient (R) was used to measure the relationship between two continuous variables. Spearman’s R correlation coefficient varies between -1 and +1 with 0 indicating no correlation. Correlations of +1 or -1 imply an exact positive or negative relationship. Continuous variables were compared between independent groups using the nonparametric Mann-Whitney U rank test. Kaplan-Meier survival curves were generated to visualize survival differences between the High-risk and Low-risk groups. The statistical significance of the survival differences was assessed using the log-rank test. Multivariable Cox proportional hazards models were used to evaluate the prognostic impact of the CPMP- derived risk group while adjusting for covariates (age, AJCC pathologic tumor stage, PR status, ER status, HER2 status, LNp). The proportional hazards assumption was checked and found to be valid for the models. Hazard ratios (HR) and 95% confidence intervals (CI) were reported. Multiple metrics, including the ROC analysis with AUC value, the PRC with the AUPRC value, the balanced accuracy, as well as the averaged Precision, Recall, and F1 score were employed to evaluate the performance of our approach and comparative methods on MP risk groups. The performance of CPMP was compared to that of comparative models and the clinical-based logistic regression model using the Wilcoxon signed-rank test on the AUROC metrics obtained from the 5-Time 5-Fold cross-validation. All tests were two-sided, and a *p*- value < 0.05 was considered statistically significant. All statistical analyses were conducted using Python 3.8 and R 4.2.2.

## Data Availability Statement

The TCGA data (slides and clinicopathological information) used in this study are available at https://portal.gdc.cancer.gov/. The follow-up data are available at TCGA Clinical Data Resource^[50]^. The TJMUCH-MP cohort is accessible upon request. The data can only be used under the condition that the request is for non-profit, purely academic research purposes, and the requesting researchers must provide valid ethics approval from their institution. The data generated in this study for the creation of the figures are provided in the Source Data file. The source codes of CPMP are available at https://github.com/lyotvincent/CPMP.

## Ethics approval

This retrospective study was approved by the Ethics Committee of Tianjin medical university cancer institute and hospital (bc20240004). Informed consent was obtained from all participants in the cohort for all data acquisition. For TCGA, no formal ethics approval was required for a retrospective study of anonymous samples.

