## Supplementary Figure S1 to S20 and Table S1 to S11. for "Predicting MammaPrint Recurrence Risk from Breast Cancer Pathological Images Using a Weakly Supervised Transformer"

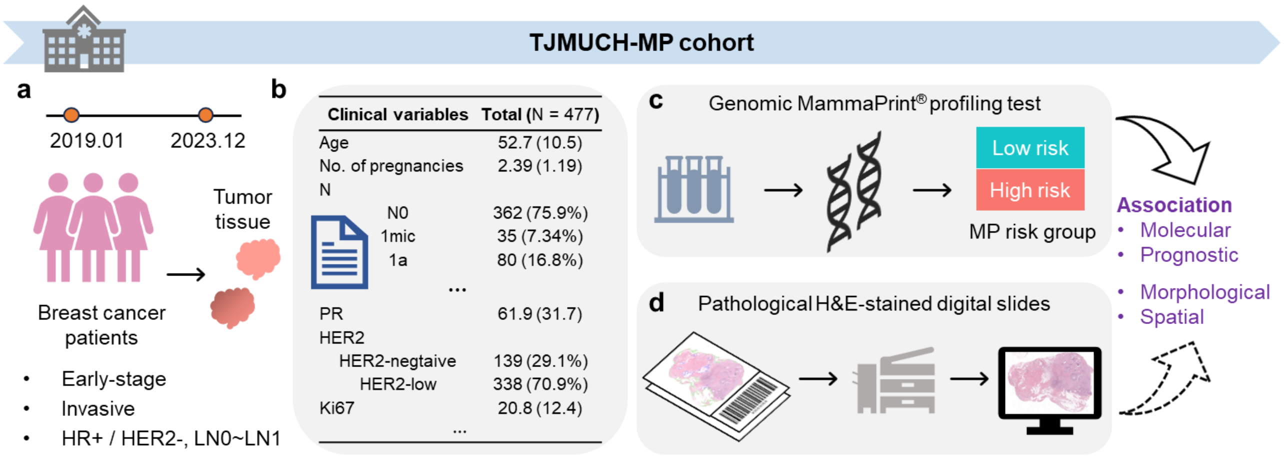


**Figure S1. Overview of the data curation criteria for the clinical TJMUCH-MP cohort.**

**a**. Schematic diagram outlining the data curation criteria for the clinical TJMUCH-MP cohort, which includes HR+/HER2- primary breast cancer patients diagnosed between January 2019 and December 2023 at Tianjin Medical University Cancer Institute and Hospital. **b**. Summary of key clinicopathologic characteristics of the cohort (N=477), including age, number of pregnancies, PR status, HER2 status, Ki-67 index, and so on. **c**. Workflow of the genomic MammaPrint (MP) profiling test, which analyzes tumor RNA to classify patients into Low-risk or High-risk groups for 10-year risk of distant metastasis. **d**. Workflow of the pathological analysis using digital whole-slide images (WSIs) of H&E-stained tissue sections. This study aims to establish an association between the molecular/prognostic information from the MP test and the morphological/spatial information derived from histopathology. HR+ =Hormone Receptor positive, HER2- =Human Epidermal growth factor Receptor 2 negative, LN0=Lymph Nodes negative, LN1=The number of Lymph Nodes positive⩽3.

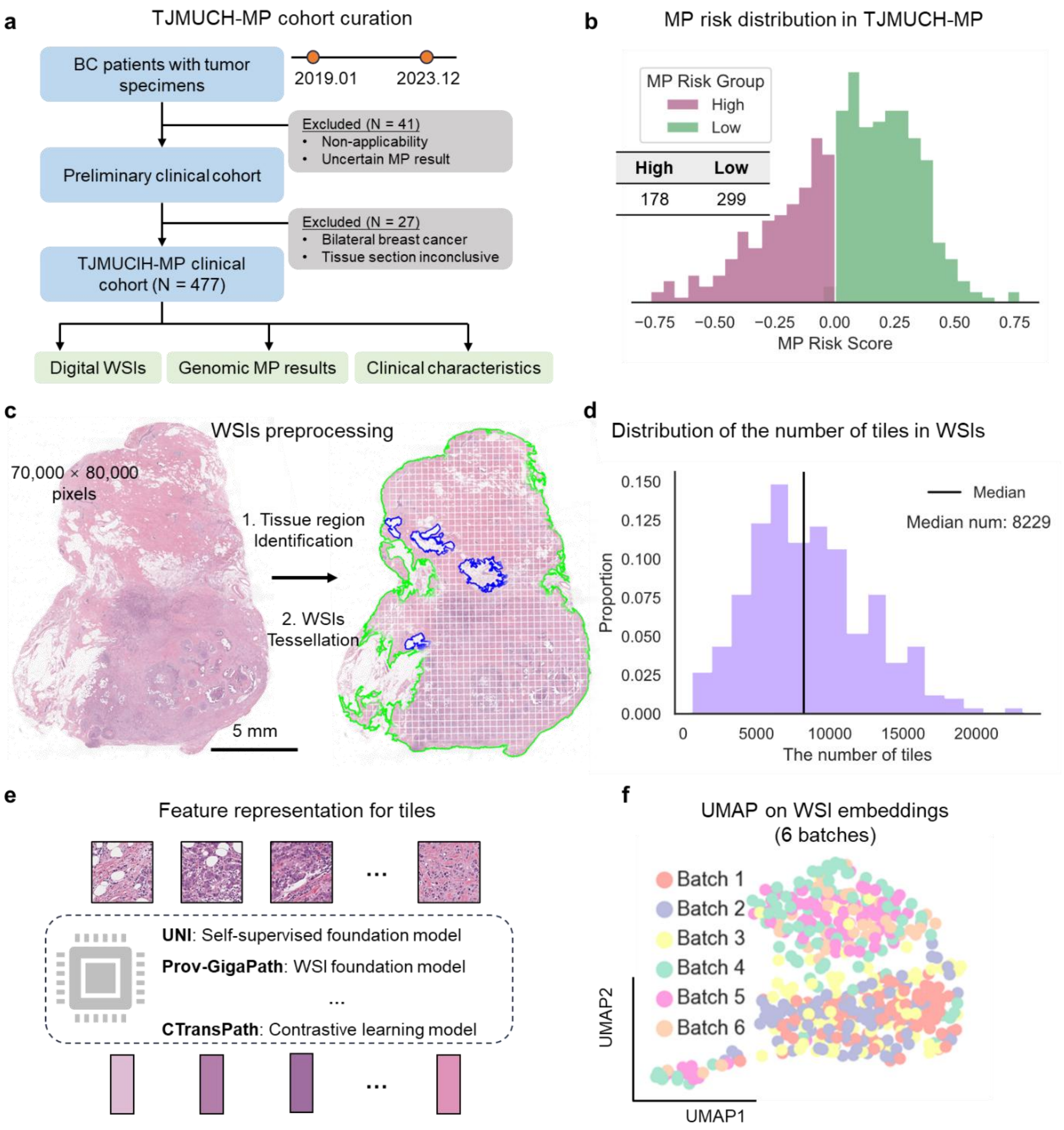


**Figure S2. Overview of the TJMUCH-MP clinical cohort and data preprocessing in CPMP.**

**a.** Data curation criteria for the TJMUCH-MP patient cohort. The HR+/HER2- BC clinical cohort was comprised of digital WSIs, MP risk results, and clinicopathologic features of 477 patients. Patients retrospectively analyzed in this cohort were diagnosed with and treated for primary BC between January 2019 and December 2023. These data were ultimately obtained for the development and evaluation of the model. **b.** Data distribution of genomic MP diagnostic results (categorized into 178 High-risk and 299 Low-risk) for patients in TJMUCH-MP cohort. **c.** Depiction of WSIs preprocessing including the foreground tissue region identification and WSIs tessellation within the tissue regions at 20× magnification. **d.** Data distribution of the number of tiles among digital WSIs in the TJMUCH-MP cohort. The median value of the number of tiles in WSIs were 8,229. **e.** Depiction of the feature extraction on tile instances for low-dimensional embedding representation. Three self-supervised foundation models including UNI, Prov-GigaPath, and CTransPath were leveraged for capturing diverse patterns from local tiles. This feature extraction part was served as the instance-level transformation function in CPMP. **f.** UMAP plot of the WSI embeddings across different scanning batches.

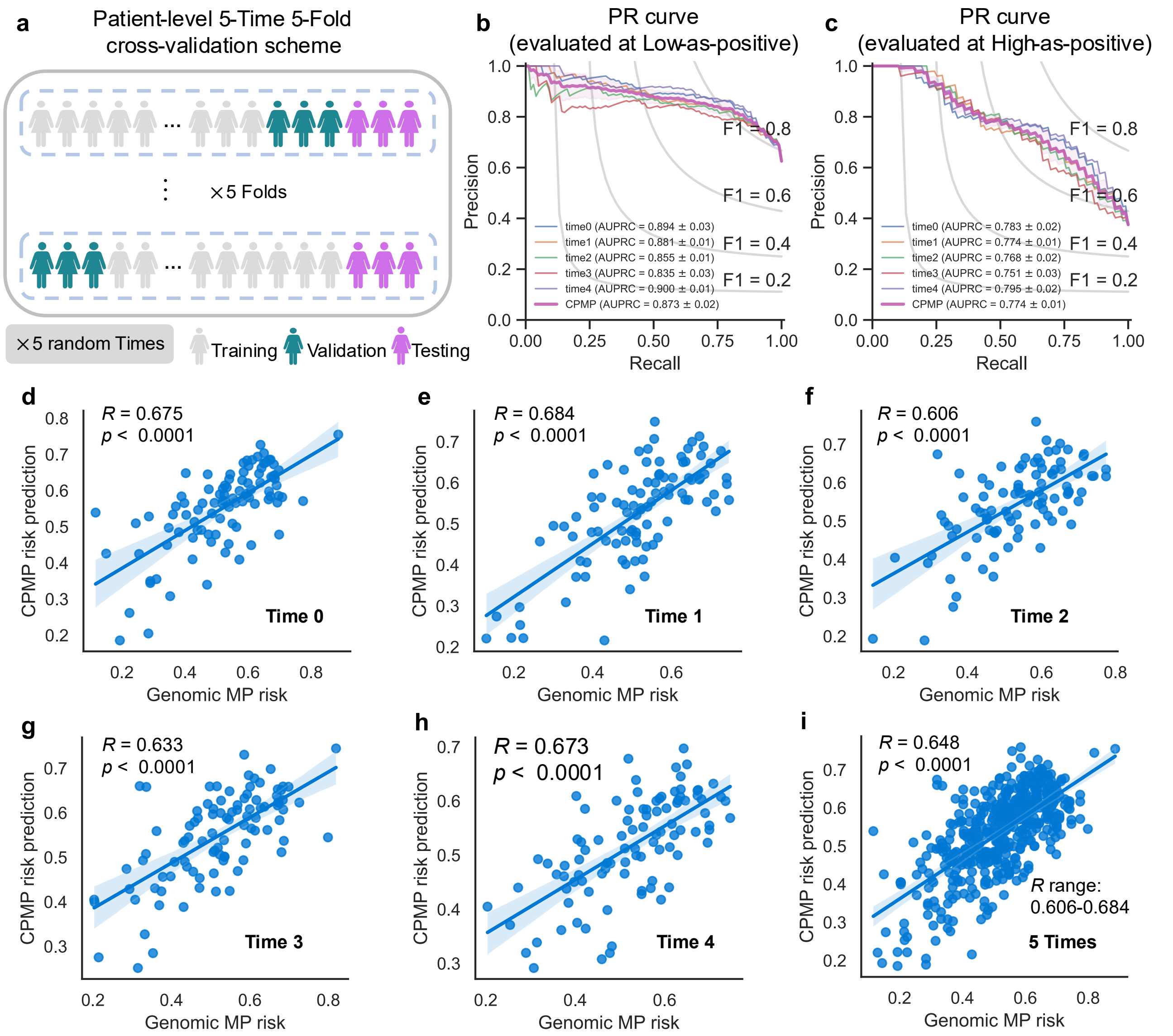


**Figure S3. Performance evaluation of the CPMP model using a patient-level 5-Time 5-Fold cross-validation strategy**

**a**. Schematic diagram illustrating the patient-level 5-Time 5-Fold cross-validation strategy employed to mitigate data bias and scanning effects. In each "Time" iteration, we randomly divided 20% of the samples for testing, while the remaining samples were evenly divided into 5 non-overlapping subsets (Folds). Slides from each patient were only assigned to one of these sets to prevent data leakage. We then iteratively trained the model on 4 subsets (4 Folds) while using the remaining subset for validation. This process was repeated for each Fold, ensuring that each Fold served as the validation subset exactly once. The trained models corresponding to the 5 Folds were evaluated on the independent testing cases. This process is repeated for 5 random iterations ("Times"), ensuring robustness. **b, c.** Precision-Recall (PR) curves for the CPMP model evaluated in both Low-as-positive (**b**) and High-as-positive (**c**) modes across the 5-Time experiments. The area under the PR curve (AUPRC) values are reported for each fold and the final CPMP model. **d-i**. Scatter plots showing the correlation between the continuous genomic MammaPrint (MP) risk score and the continuous CPMP-predicted risk probability for each patient in the test set of 5-Time experiments (from Time-0 to Time-4, **d-e**) and the combined results from all five times (**i**). The Spearman correlation coefficient (R) and p-value are calculated for each of them. The fitted linear regression line and its 95% confidence interval are also shown. The range of R values across the 5-Time experiments is indicated in panel **i**.

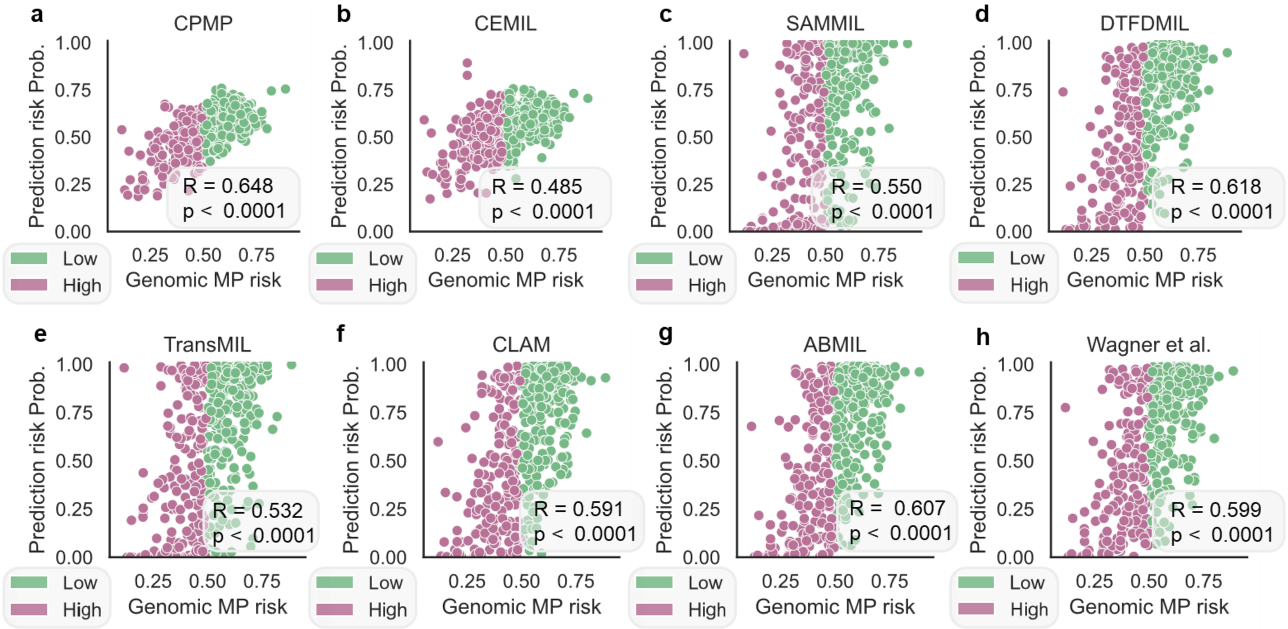


**Figure S4. Performance comparison of CPMP with state-of-the-art weakly supervised methods in predicting MammaPrint recurrence risk.**

**a-h**. Scatter plots showing the correlation between the continuous genomic MP risk score and the predicted risk probability for each patient in the test set, using the proposed CPMP model (**a**) and seven comparative methods: CEMIL (**b**), SAMMIL (**c**), DTFDMIL (**d**), TransMIL (**e**), CLAM (**f**), ABMIL (**g**), and Wagner et al. (**h**). The data points are colored by the corresponding genomic MP risk group (Low-risk in Spring green, High-risk in Lilac). The Spearman correlation coefficient (R) and p-value are reported for each method. All models were evaluated using the same patient-level 5-Time 5-Fold cross-validation strategy on the TJMUCH-MP cohort. The results demonstrate that CPMP achieves the highest correlation (R = 0.648) with the genomic MP risk scores, outperforming all other comparative methods.

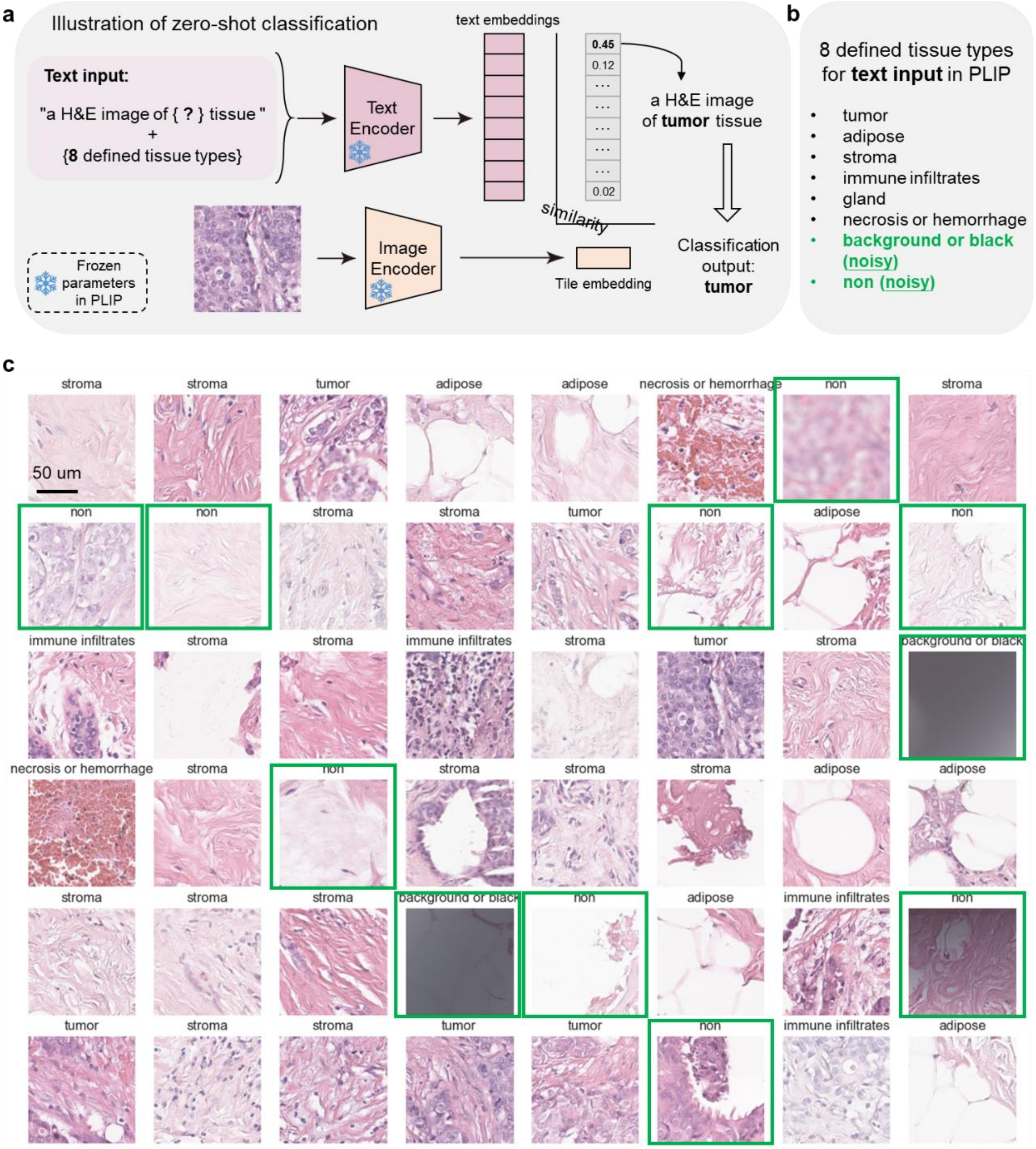


**Figure S5. Depiction of the zero-shot classification experiment for noisy tissue-type filtering.**

**a.** Illustration of zero-shot tissue type classification. All tiles were determined as one of 8 tissue types by performing zero-shot classification using the visual-language foundation PLIP model. **b**. 8 defined tissue types: tumor, adipose, stroma, immune infiltrates, gland, and necrosis or hemorrhage, background or black, and ‘non’, which were served as text input for the PLIP model. **c**. Zero-shot classification results of tiles. Tiles in green box categorized as ‘background or black’ or ‘non’ tissue type were considered as noisy tiles and filtered out. Tiles identified as specific 6 tissue types (tumor, adipose, stroma, immune infiltrates, gland, and necrosis or hemorrhage) were then retained for the comparative model. AUROC=Area Under the Receiver Operating Characteristic, w/=With, w/o=Without, ns=Statistically no significance.

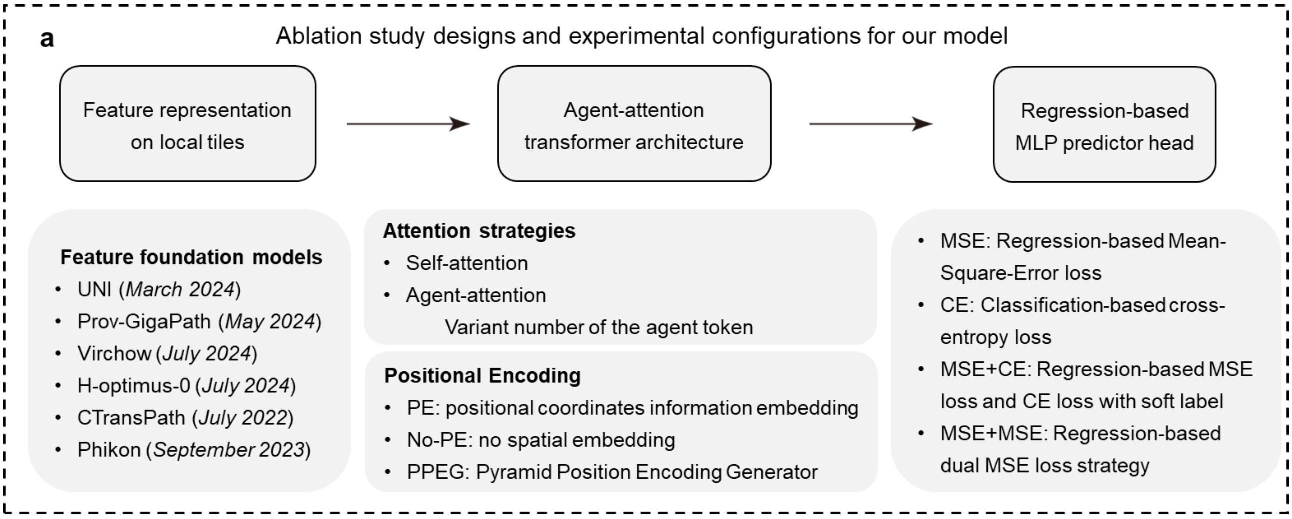


**Figure S6. Illustration of experimental designs in the ablation study.**

**a**. The design outlines of the ablation study and the experimental configurations utilized for the ablation models. The ablation study focused on evaluating various foundational models for feature representation, as well as different attention strategies and positional encoding methods within the transformer architecture. Additionally, we examined various loss strategies in the MLP predictor head.

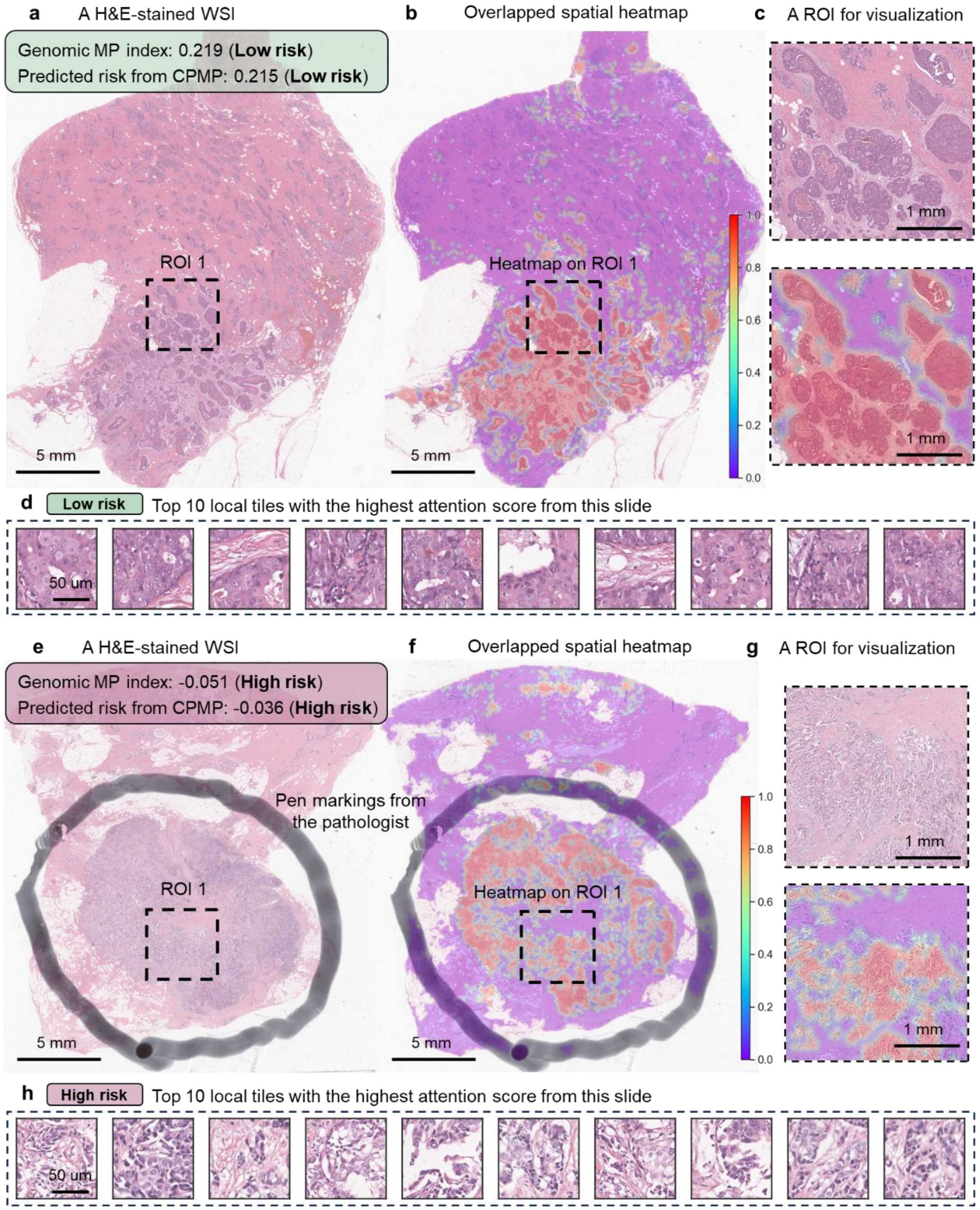


**Figure S7. Whole-slide-level spatial heatmap visualization illustrating the spatial arrangement and morphological patterns of tumors.**

**a, e**. Example H&E-stained WSIs of patients with genomic MP Low-risk (**a**) and High-risk (**e**) in the TJMUCH-MP clinical cohort. Genomic MP diagnostic risk index and corresponding prediction risk results from CPMP were marked in the box. Scale bar: 5 mm / 3 mm. **b, f**. The overlapped spatial heatmap visualization for the corresponding Low-risk (**b**) and High-risk (**f**) cases at the whole-slide-level. The tile-level attention scores that contributed to the final patient-level risk prediction were yielded on the agent-attention matrix between local tiles using the attention rollout method. Spatial heatmap results were then generated by mapping the attention scores of all local tiles to their corresponding locations within the WSIs and they were superposed onto the histopathological WSIs for interpretability. Please see Methods section. A colorbar was added on the lower right of the heatmap. Scale bar: 5 mm / 3 mm. **c, g.** Fine-grained pathological ROIs extracted from the WSIs and the corresponding spatial heatmap visualization results. Scale bar: 500 μm. **d, h.** Top 10 local tiles with the highest attention score from each slide for both MP Low-risk (**d**) and High-risk (**h**) cases. Scale bar: 50 μm. WSI=Whole Slide Image, ROI=Region of Interest, MP=MammaPrint diagnostic test.

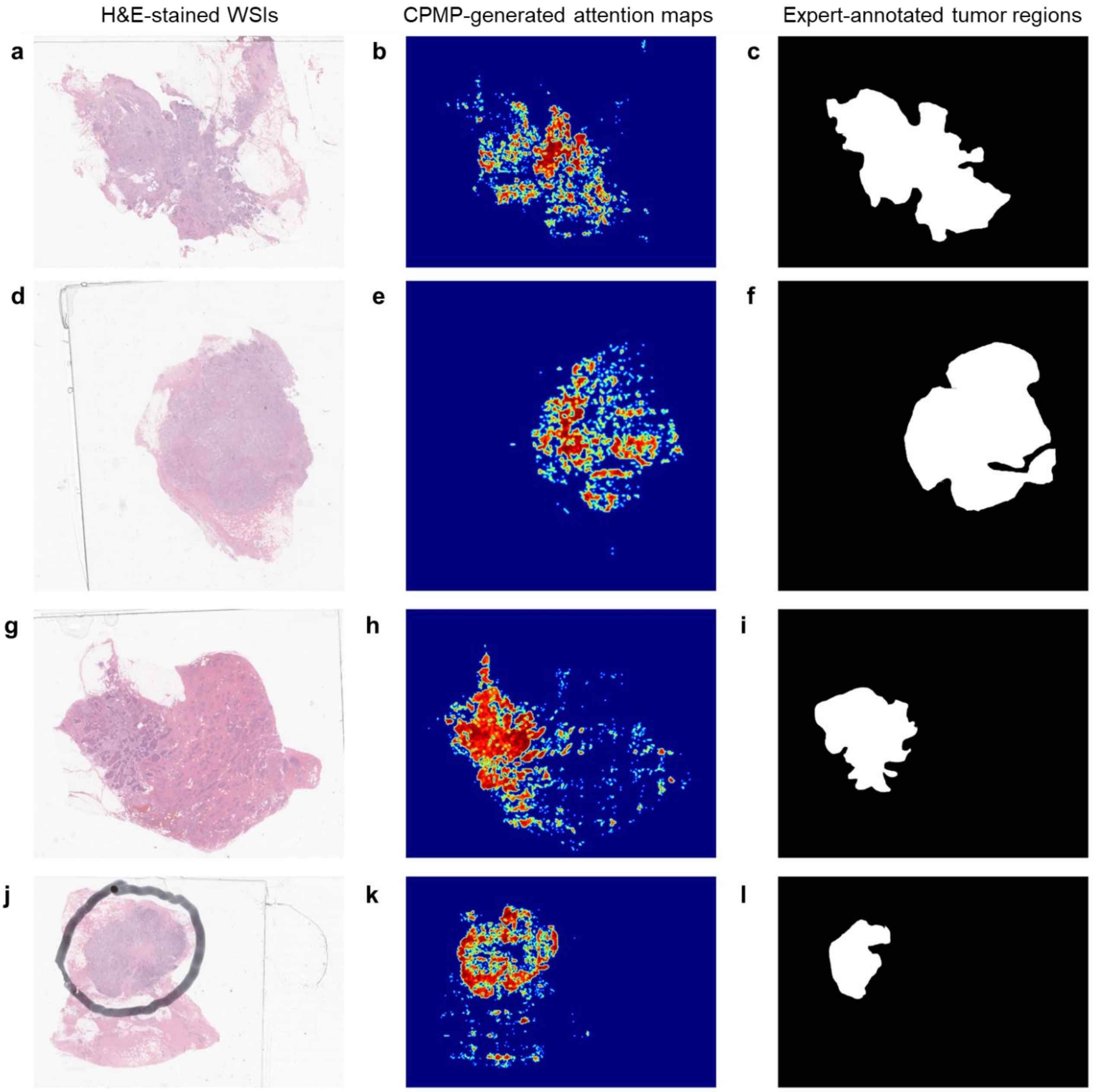


**Figure S8. Visual comparison of CPMP-generated attention maps with expert-annotated tumor regions across representative histopathology slides.**

**a, d, g, j**. Original H&E-stained whole slide images showing diverse tissue morphologies and tumor architectures. **b, e, h, k.** Corresponding attention heatmaps generated by CPMP, highlighting spatially resolved predictive scores (red = high; blue = low) for tumor-related features. **c, f, i, l.** Binary tissue masks of manually annotated tumor areas from pathologists. Panels **a–c**, **d–f**, **g–i**, and **j–l** represent four distinct cases, illustrating the consistency across diverse samples.

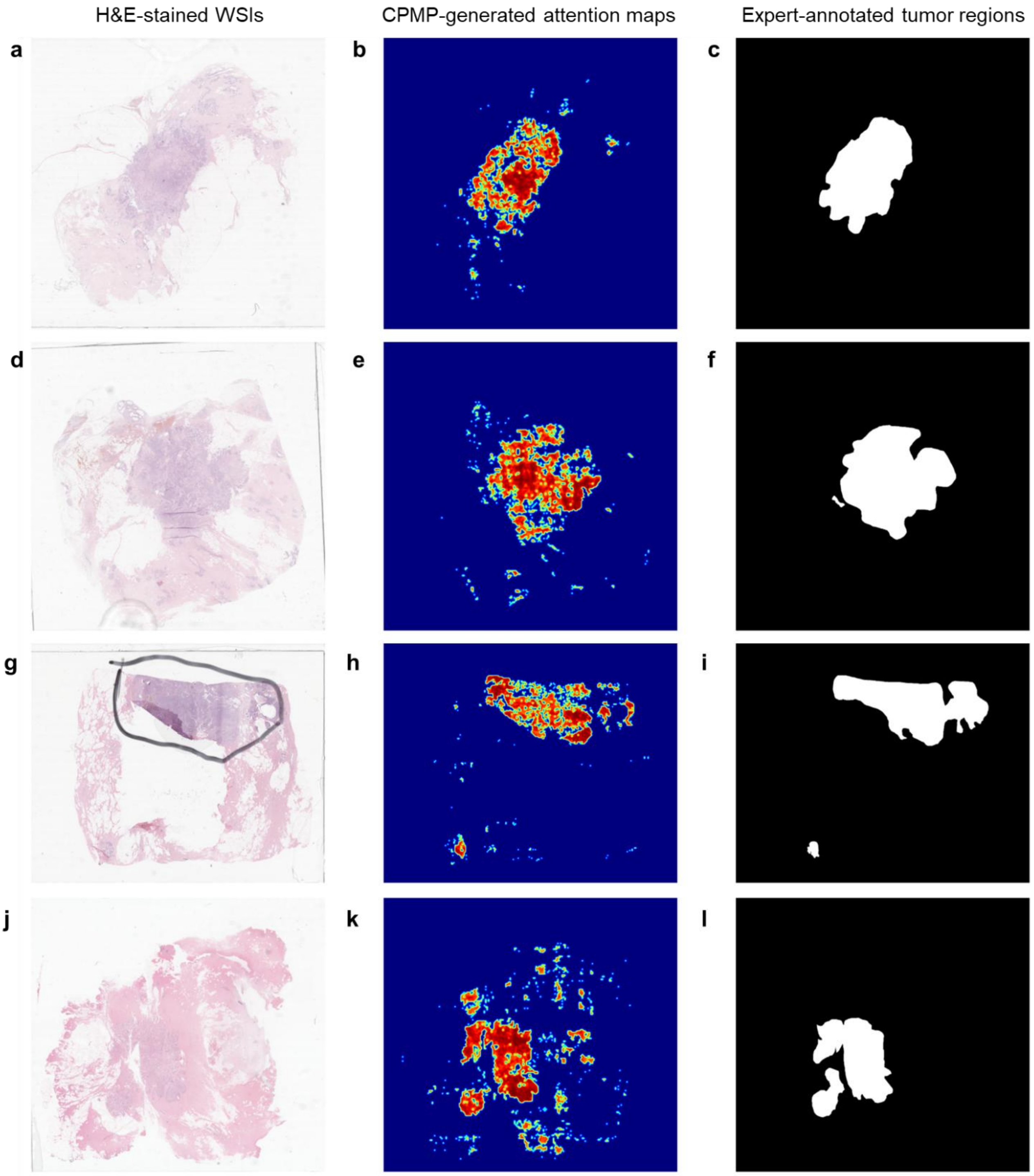


**Figure S9. Visual comparison of CPMP-generated attention maps with expert-annotated tumor regions across representative histopathology slides.**

**a, d, g, j**. Original H&E-stained whole slide images showing diverse tissue morphologies and tumor architectures. **b, e, h, k.** Corresponding attention heatmaps generated by CPMP, highlighting spatially resolved predictive scores (red = high; blue = low) for tumor-related features. **c, f, i, l.** Binary tissue masks of manually annotated tumor areas from pathologists. Panels **a–c**, **d–f**, **g–i**, and **j–l** represent four distinct cases, illustrating the consistency across diverse samples.

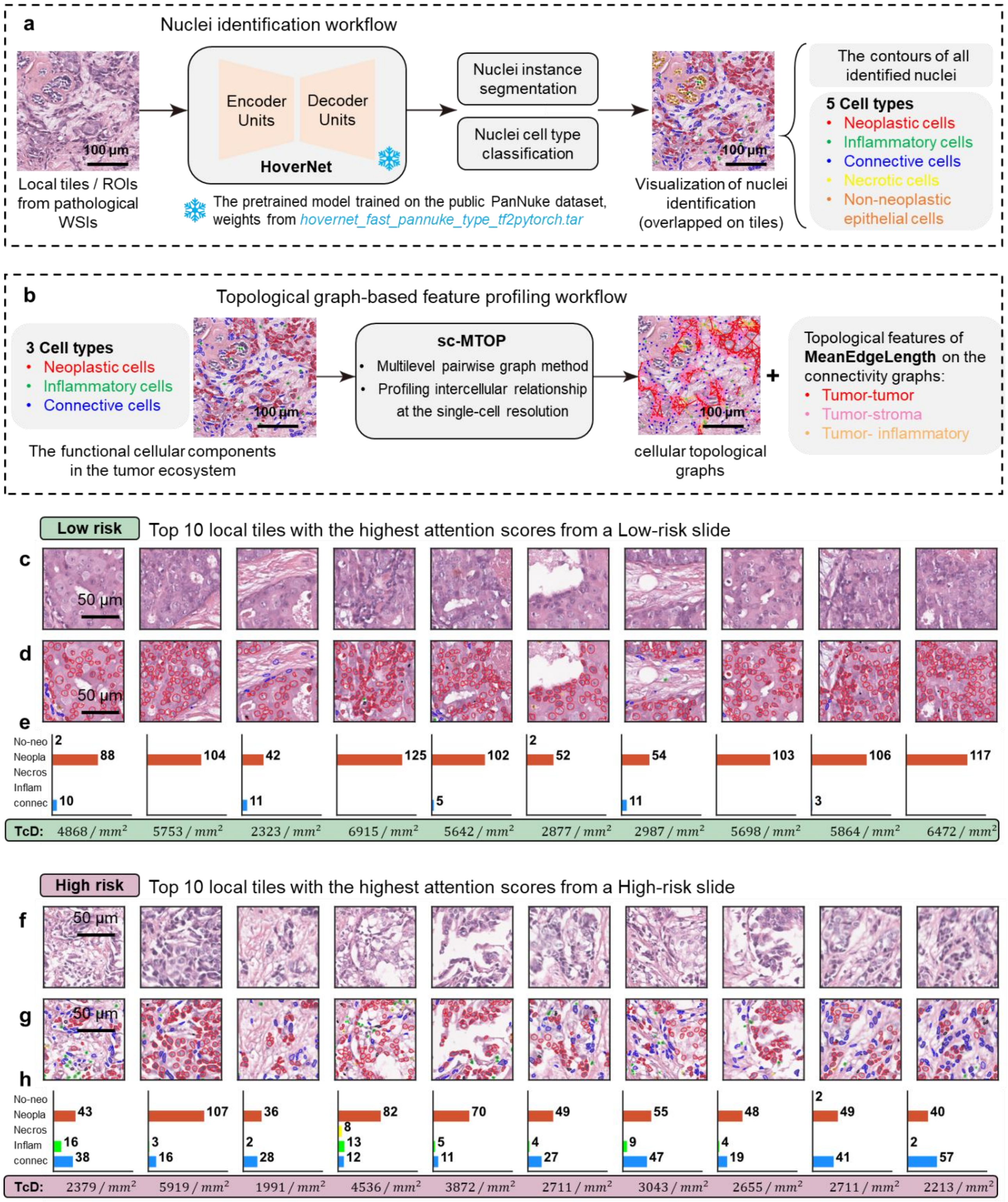


**Figure S10. Illustration of nuclei identification and intercellular connectivity graph profiling workflow.**

**a**. Illustration of the nuclei identification workflow using HoverNet. HoverNet method was applied to local tiles or ROIs for nuclei simultaneous instance segmentation and cell type classification. The pretrained model trained on the public PanNuke dataset was utilized, and the model weights were loaded from hovernet_fast_pannuke_type_tf2pytorch.tar. Each cell nucleus was classified as one of five cell types (neoplastic, inflammatory, connective, necrotic, and non-neoplastic epithelial cells). The contours of identified nuclei were located and drawn on the fine-grained pathological ROIs. **b.** Illustration of the topological graph-based feature profiling workflow using sc-MTOP framework. Three main cellular components (neoplastic, inflammatory, and connective stromal cells) in the tumor ecosystem were employed to construct topological connectivity graphs among these detected cell nuclei. The sc-MTOP framework was leveraged for extracting the graph-based features at the single-cell level. The topological features of graph-edge length-related MeanEdgeLength were yielded to characterize the intercellular spatial relationships within tumors. The MeanEdgeLength metric reflects the average distance of a cell interacting with its neighboring cells. It was calculated using the connectivity graphs of tumor-tumor, tumor-stroma, and tumor-inflammatory cell interactions for each tumor cell. **c, d, f, g**. Nuclei segmentation results for top 10 local tiles with the highest attention score from the corresponding MP Low-risk (**c**) and High-risk (**f**) slides. Scale bar: 50 μm. **e, h**. The cellular composition quantitation of the five cell types in each local tile region. The number of cells for each type were labelled on the right side of the plot and the TcD values representing the density of tumor cells were marked in the lower part of the plot. WSI=Whole Slide Image, ROI=Region of Interest, TcD=Tumor cells Density.

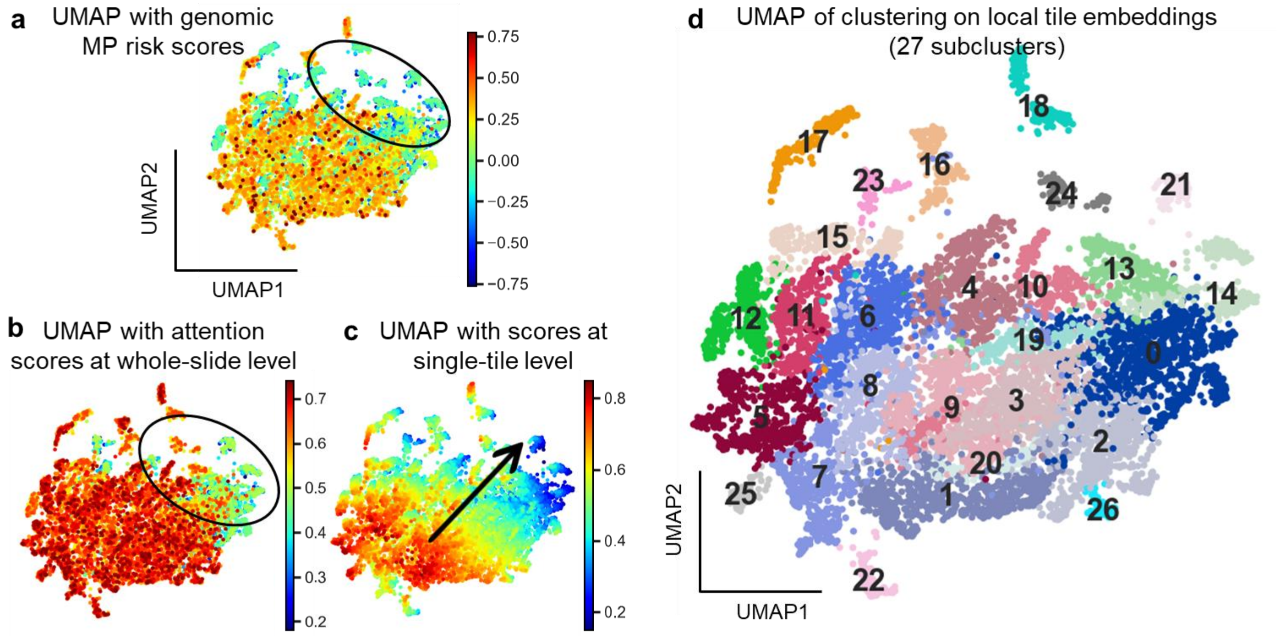


**Figure S11. Characterization of MP-associated tumor morphological phenotypes discovered by CPMP.**

**a-c**. UMAP plots of learned embeddings of local tiles sampled from WSIs in the test set, colored by genomic MP risk scores (**a**), local attention scores from CPMP at the whole-slide level (**b**), and prediction probability scores from CPMP at the single-tile level (**c**). For each WSI, top 100 tiles with the highest attention scores were identified as representative local regions to create a collection of tiles. Each tile in this collection was then fed into CPMP for feature embedding extraction. The arrow in panel **c** indicates the trajectory direction of recurrence risk development. **d**. UMAP plot of unsupervised clustering results of learned embeddings of local tiles. Clusters are labeled with inferred subcluster IDs (subclusters were named as SC0-SC26 and assigned colors for visualization).

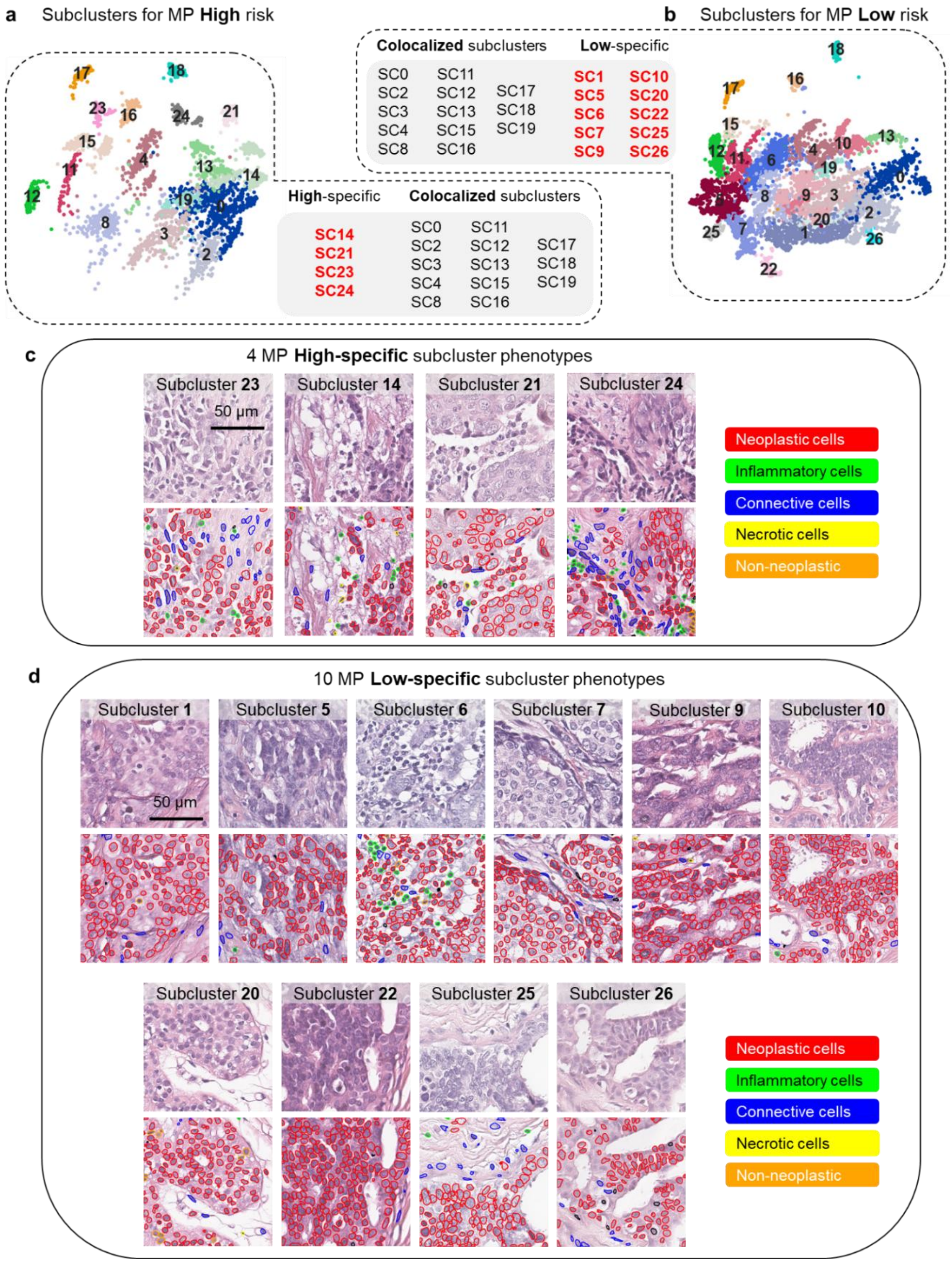


**Figure S12. The characterization of MP-associated tumor morphological phenotypes discovered by CPMP.**
**a, b.** UMAP subplots of tile embeddings for subclusters associated with MP High (**a**) and Low (**b**) risk. The IDs of High-specific, Low-specific, and Colocalized subclusters were also listed. **c,d.** Depiction of MP risk-specific subcluster phenotypes. 4 MP-High-specific and 10 MP-Low-specific subclusters were identified. The representative tiles and the corresponding nuclei segmentation results in High-specific (**c**) and Low-specific (**d**) subclusters demonstrated the phenotypic characteristics of different MP risk groups. HoverNet method was applied for nuclei simultaneous segmentation and classification.

**a**

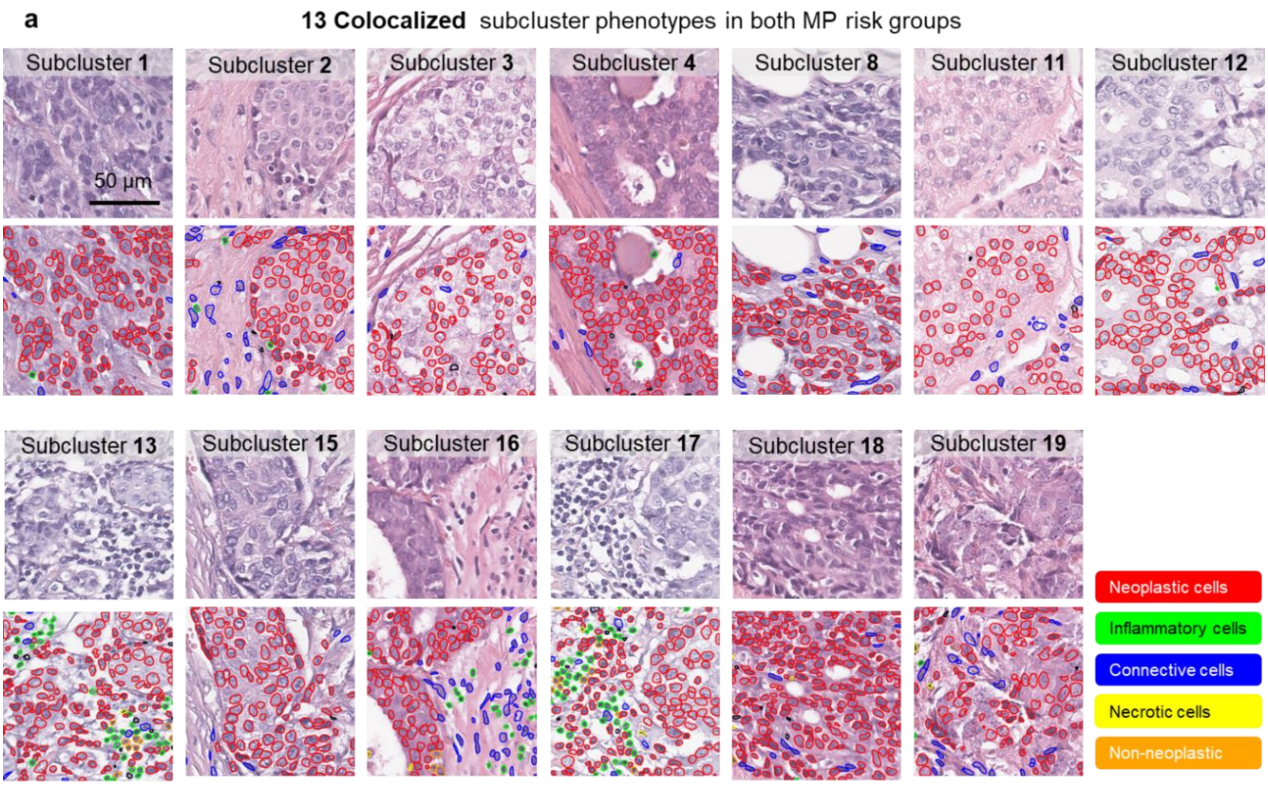


**b** Quantitative results of cellular composition of 5 cell types among 27 subclusters

| Cell types | Subcluster ID | | | | | | | | | | | | |
|---|---|---|---|---|---|---|---|---|---|---|---|---|---|
| | SC13 | SC0 | **SC21** | **SC23** | **SC14** | **SC24** | SC19 | SC4 | SC15 | SC8 | SC16 | SC18 | SC3 | SC17 |
| connec | 15.71 | 16.59 | 15.41 | 18.98 | 17.45 | 12.86 | 10.96 | 16.37 | 15.57 | 15.52 | 15.00 | 15.26 | 14.86 | 14.50 |
| inflam | 4.76 | 4.37 | 3.07 | 3.56 | 3.72 | 2.80 | 1.64 | 2.50 | 3.08 | 2.86 | 1.54 | 2.54 | 2.17 | 2.39 |
| necros | 0.76 | 0.65 | 0.81 | 0.73 | 0.72 | 0.33 | 0.38 | 0.49 | 0.52 | 0.46 | 0.38 | 0.41 | 0.37 | 0.37 |
| neopla | 59.66 | 55.84 | 60.97 | 57.08 | 62.38 | 68.71 | 64.01 | 60.26 | 61.93 | 63.22 | 64.69 | 66.04 | 64.17 | 65.43 |
| no-neo | 0.67 | 0.31 | 0.21 | 0.18 | 0.44 | 1.30 | 0.23 | 0.58 | 0.46 | 0.62 | 0.36 | 0.47 | 0.48 | 0.67 |

| Cell types | Subcluster ID | | | | | | | | | | | | |
|---|---|---|---|---|---|---|---|---|---|---|---|---|---|
| | **SC26** | SC25 | **SC5** | SC12 | **SC22** | **SC9** | **SC20** | SC2 | **SC6** | SC11 | **SC10** | **SC1** | **SC7** |
| connec | 16.74 | 16.65 | 18.04 | 14.81 | 15.58 | 16.59 | 16.99 | 18.04 | 15.81 | 17.11 | 16.80 | 16.36 | 16.85 |
| inflam | 2.10 | 2.57 | 1.90 | 1.77 | 1.81 | 1.62 | 1.90 | 2.67 | 1.78 | 1.90 | 2.14 | 1.88 | 2.19 |
| necros | 0.67 | 0.35 | 0.29 | 0.31 | 0.29 | 0.34 | 0.30 | 0.44 | 0.44 | 0.44 | 0.45 | 0.38 | 0.38 |
| neopla | 55.64 | 50.92 | 56.96 | 59.93 | 57.61 | 57.06 | 59.01 | 57.78 | 57.98 | 59.20 | 59.03 | 59.82 | 60.04 |
| no-neo | 1.76 | 0.22 | 0.59 | 0.52 | 0.31 | 0.38 | 0.36 | 0.48 | 0.44 | 0.59 | 0.71 | 0.50 | 0.61 |

**High-specific** **Low-specific** **Colocalized**

**Figure S13. The characterization of morphological phenotypes discovered by CPMP.**

**a**. Depiction of morphological phenotypes of 13 colocalized subclusters presented in both MP risk groups. The representative tiles and the corresponding nuclei segmentation results in colocalized subclusters reflected the consistent morphological characteristics of early-stage BC patients to whom the genomic MP test is applicable. **b**. The quantitative cellular composition results of 27 subclusters in terms of the five cell types (connective, inflammatory, necrotic, neoplastic, and non-neoplastic epithelial cells). The Subclusters were sorted according to the hierarchical clustering results in Figure 5k. Distinct subcluster types (High-specific, Low-specific, and Colocalized subclusters) were highlighted in different colors.

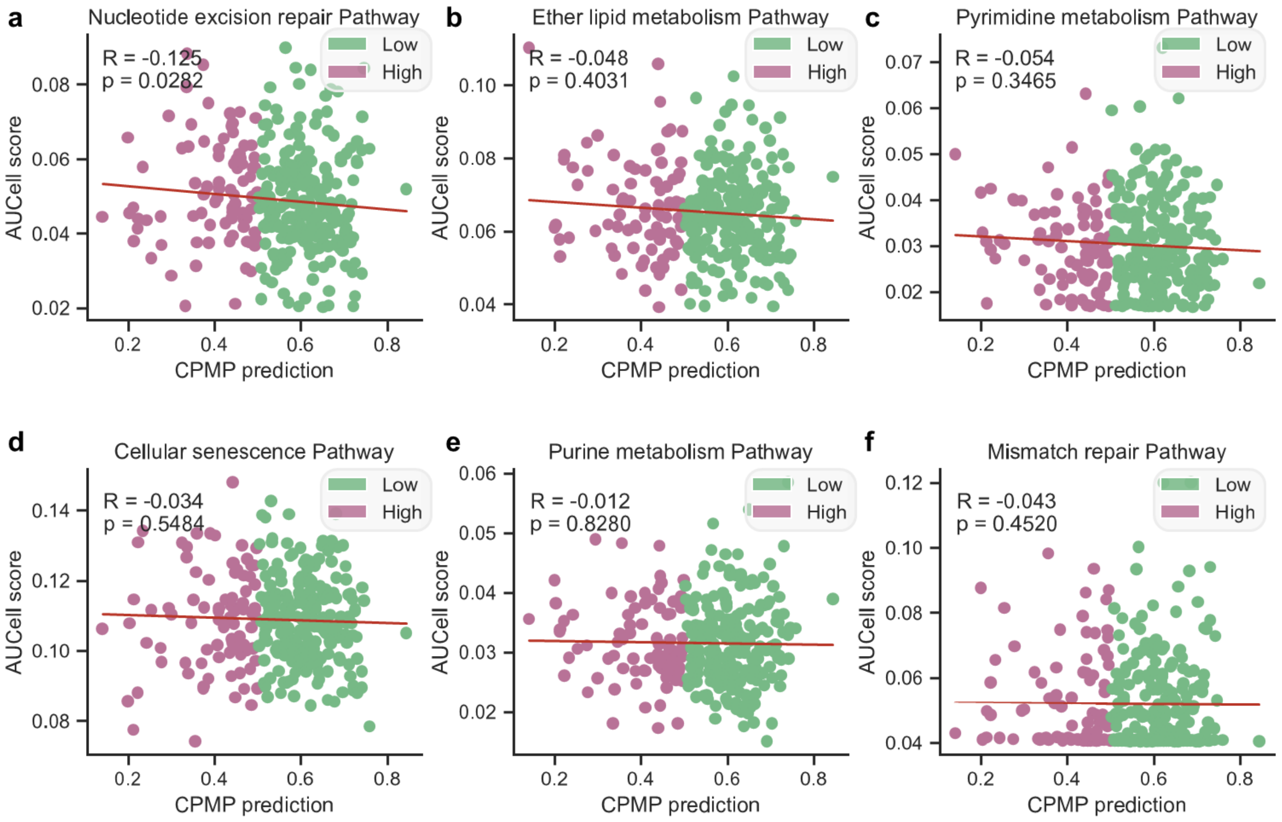


**Figure S14. Correlation between CPMP-predicted risk and gene set activity scores for key biological pathways.** **a-f**. Scatter plots showing the correlation between the continuous CPMP-predicted risk score (x-axis) and the AUCell score (y-axis) of six key biological pathways derived from the MammaPrint (MP) gene signature: (**a**) Nucleotide excision repair, (**b**) Ether lipid metabolism, (**c**) Pyrimidine metabolism, (**d**) Cellular senescence, (**e**) Purine metabolism, and (**f**) Mismatch repair. The data points are colored by the corresponding genomic MP risk group (Low-risk in Spring green, High-risk in Lilac). The Spearman correlation coefficient (R) and p-value are reported for each pathway. The fitted linear regression line is also shown. All analyses were performed on the TJMUCH-MP cohort. No significant correlations were found for these specific pathways.

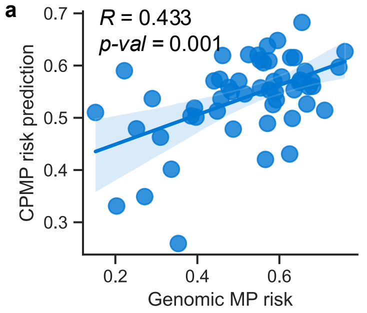


b Performance evaluation of CPMP in the external cohort (n=54)

| Performance | Evaluation metrics | | | | | | |
|---|---|---|---|---|---|---|---|
| | **AUROC** | **High (n=20) F1-score** | **Low (n=34) F1-score** | **Weighted avg. precision** | **Weighted avg. recall** | **Weighted avg. F1-score** | **Balanced accuracy** |
| CPMP | **0.772** | 0.651 | 0.769 | 0.733 | 0.722 | 0.726 | 0.718 |

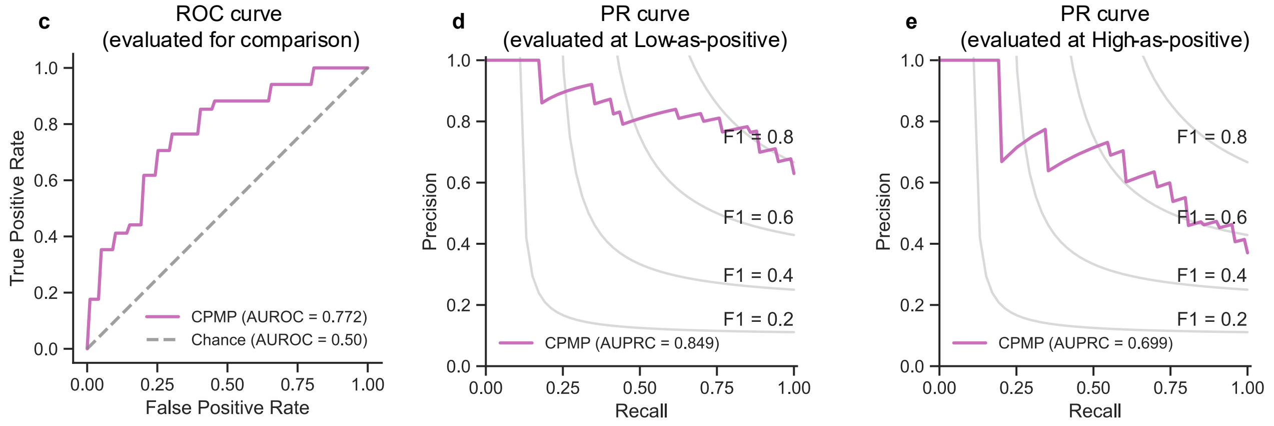


**Figure S15. Performance evaluation for MP recurrence risk prediction in the external cohort.**

**a**. Scatter plot showing the correlation between the continuous genomic MP risk score and the continuous CPMP-predicted risk probability for each patient in the external cohort (n=54), with a Spearman correlation coefficient (R) of 0.433 (p-value = 0.001). The fitted linear regression line is also shown. **b**. Performance evaluation results of CPMP in the external cohort (n=54). **c-e**. External evaluation of CPMP for MP recurrence risk group prediction using multiple metrics: (**c**) ROC curve, (**d**) Precision-Recall (PR) curve at the Low-as-positive mode, and (**e**) PR curves at the High-as-positive mode. Low-as-positive is the condition that we aim to focus on Low-risk patients to avoid overtreatment. High-as-positive is to observe the High-risk patients who may be at risk of recurrence.

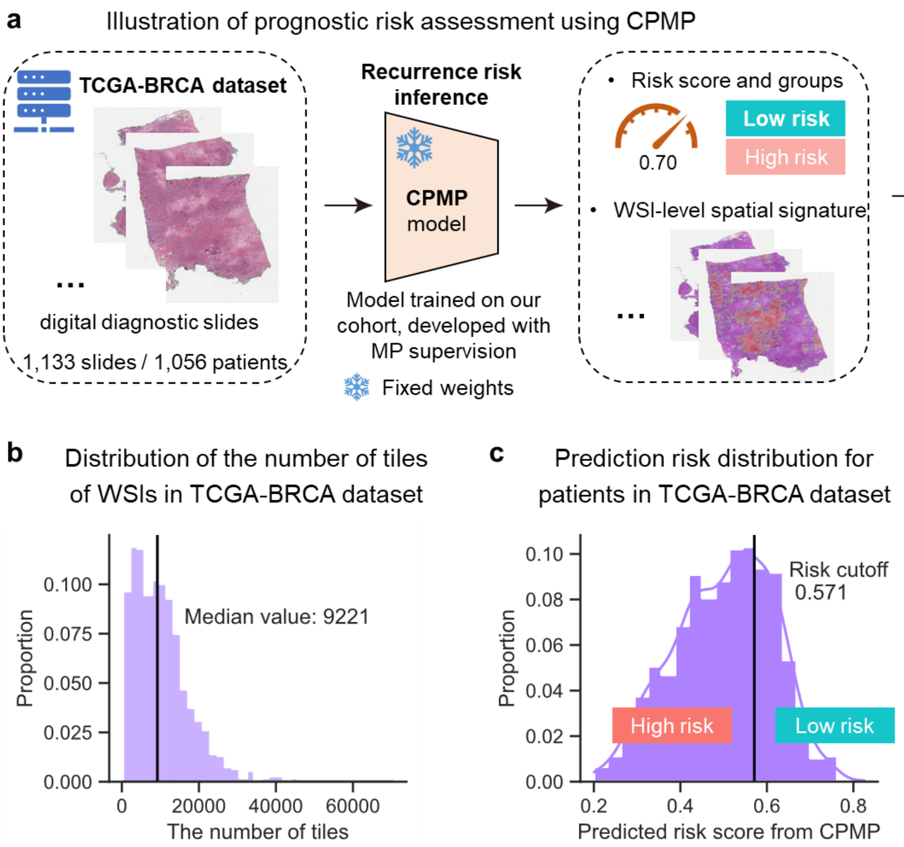


**Figure S16. Prognostic risk assessment workflow on external TCGA-BRCA data using CPMP.**

**a**. Illustration of prognostic risk assessment workflow using CPMP. CPMP model was applied to the TCGA-BRCA cohort, which includes 1,056 patients with digital diagnostic slides as well as clinical follow-up information. All digital slides in the TCGA-BRCA cohort were inferred by CPMP trained on the TJMUCH-MP cohort to predict recurrence risk scores. The prediction results of multiple slides from the same patient were averaged into the patient-level risk values. The continuous risk score at the patient-level was then binarized into High-risk and Low-risk groups by the cutoff value. The prognostic analysis was performed in terms of Distant metastasis-free Survival (DMFS), Disease-free Interval (DFI), Progression-free Interval (PFI), and Overall Survival (OS) indicators. **b**. The distribution of the number of tiles of digital WSIs in the TCGA-BRCA dataset. The median value of the number of tiles in diagnostic slides were 9,221. **c**. Data distribution of predicted risk probabilities for patients in the TCGA-BRCA dataset. Patients were categorized into High-risk and Low-risk groups based on a cutoff value determined by the corresponding CPMP model. TCGA=The Cancer Genome Atlas, MP=MammaPrint diagnostic test, WSI=Whole Slide Image.

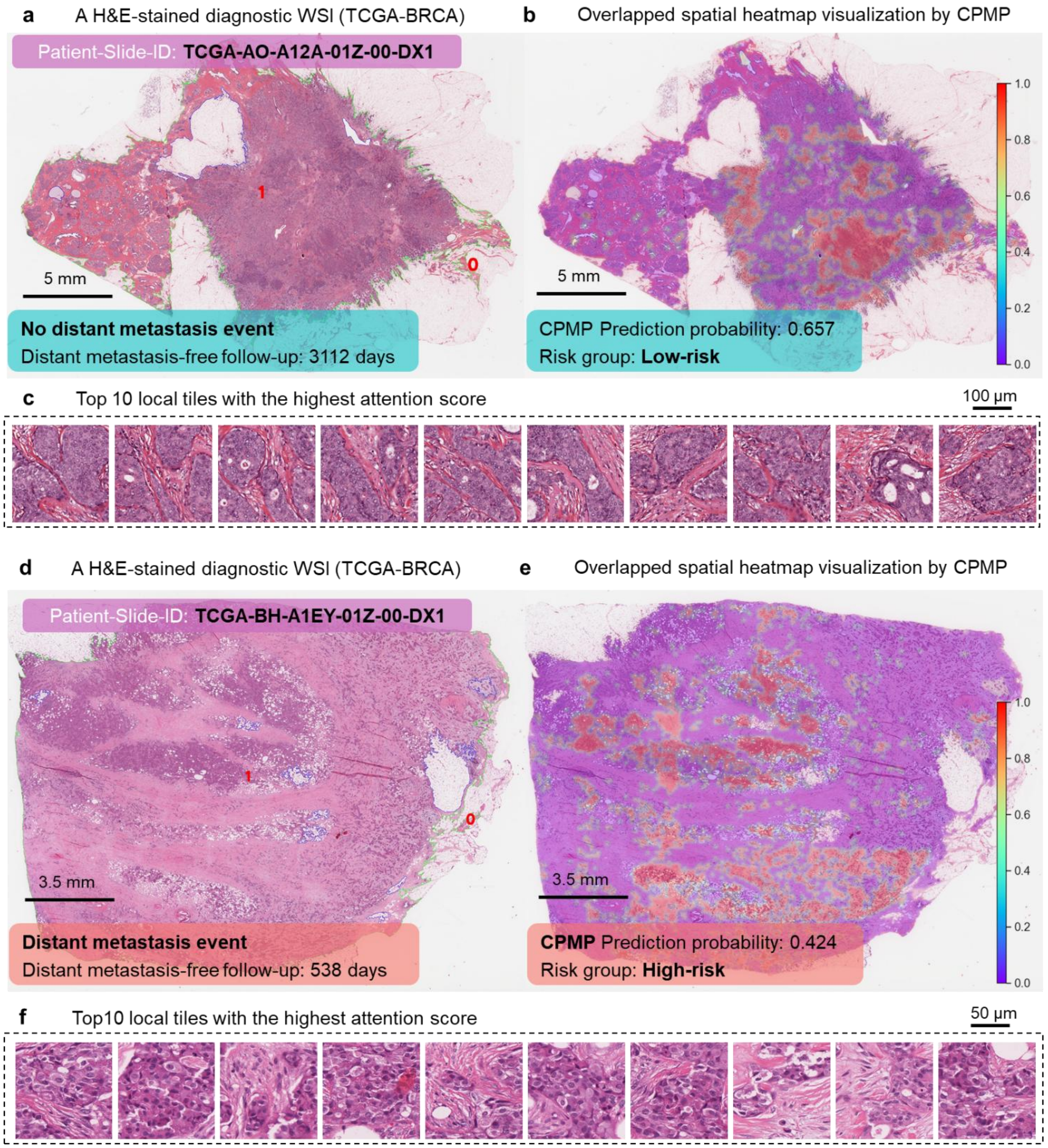


**Figure S17. Whole-slide-level spatial heatmap visualization for morphological patterns associated with prognostic events on the external TCGA-BRCA data.**

**a**. H&E-stained diagnostic WSIs of patients with no distant metastasis event in the TCGA-BRCA dataset. Scale bar: 5 mm. **b**. The whole-slide-level overlapped spatial heatmap visualization derived from CPMP. The tile-level attention scores that contributed to the final patient-level risk prediction were yielded on the agent-attention matrix between local tiles using the attention rollout method. Spatial heatmap results were then generated by mapping the attention scores of all local tiles to their corresponding locations within the WSIs and they were superposed onto the histopathological WSIs for interpretability. Scale bar: 5 mm. A colorbar was added on the right of the heatmap. The patient ID, distant metastasis-free follow-up period, and prediction risk probability from CPMP were marked in the boxes. The slide was predicted as

Low-risk group by CPMP. **c**. Top 10 local tiles with the highest attention score revealing the tumor morphological patterns of the slide. Scale bar: 100 µm. **d**. H&E-stained diagnostic WSIs of patients with distant metastasis event in the TCGA-BRCA dataset. Scale bar: 3.5 mm. **e**. The whole-slide-level overlapped spatial heatmap visualization derived from CPMP. Scale bar: 3.5 mm. The patient ID, distant metastasis-free follow-up period, and prediction risk probability from CPMP were marked in the boxes. This slide was predicted as High-risk group by CPMP. **f**. Top 10 local tiles with the highest attention score revealing the tumor morphological patterns of the slide. Scale bar: 50 µm. WSI=Whole Slide Image.

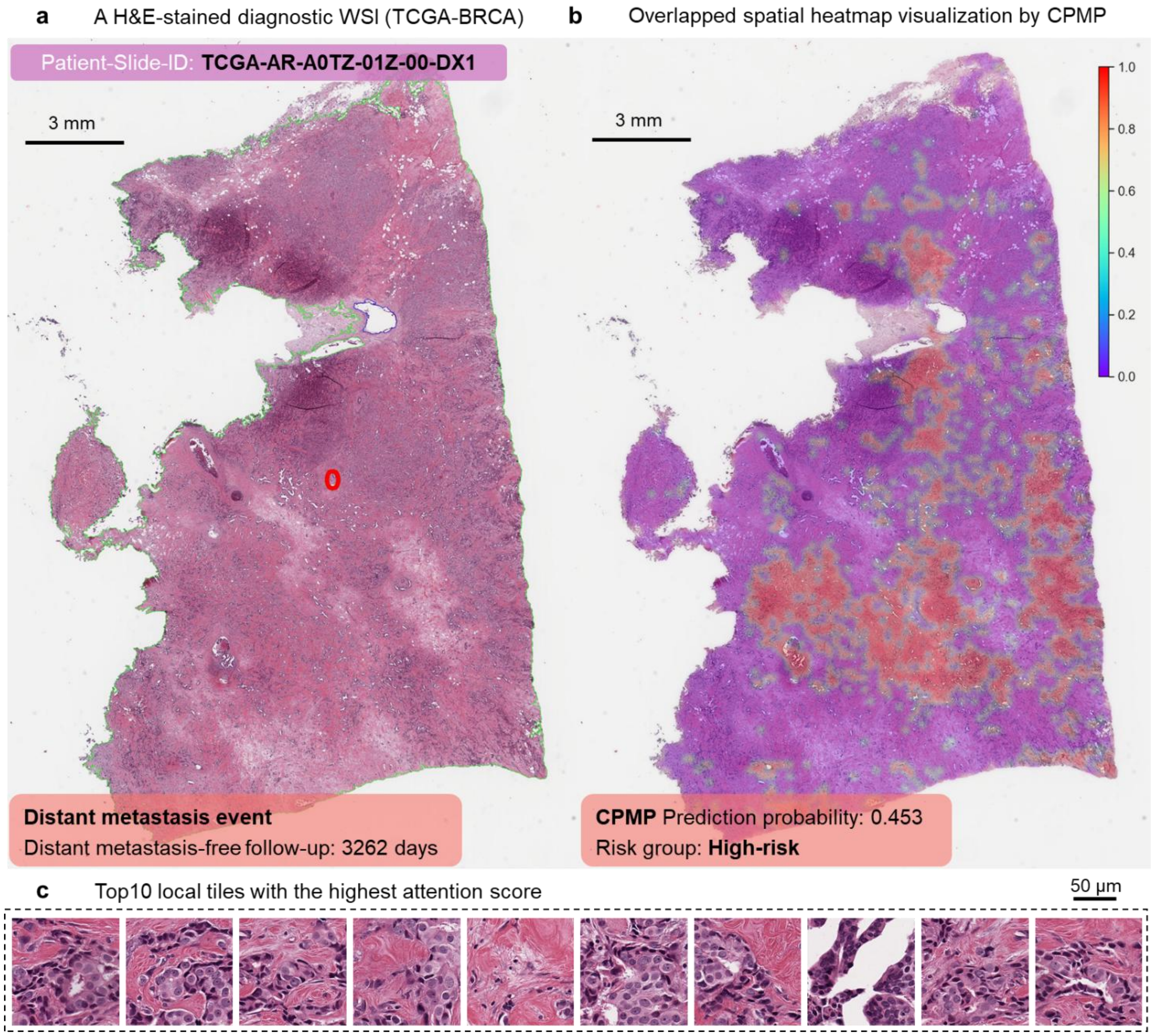


**Figure S18. Whole-slide-level spatial heatmap visualization for morphological patterns associated with distant metastasis event on the external TCGA-BRCA data.**

**a.** H&E-stained diagnostic WSIs of patients with distant metastasis event in the TCGA-BRCA dataset. Scale bar: 3 mm. **b**. The whole-slide-level overlapped spatial heatmap visualization derived from CPMP. Scale bar: 3 mm. The patient ID, distant metastasis-free follow-up period, and prediction risk probability from CPMP were marked in the boxes. This slide was predicted as High-risk group by CPMP. **f**. Top 10 local tiles with the highest attention score revealing the tumor morphological patterns of the slide. Scale bar: 50 µm. WSI=Whole Slide Image.

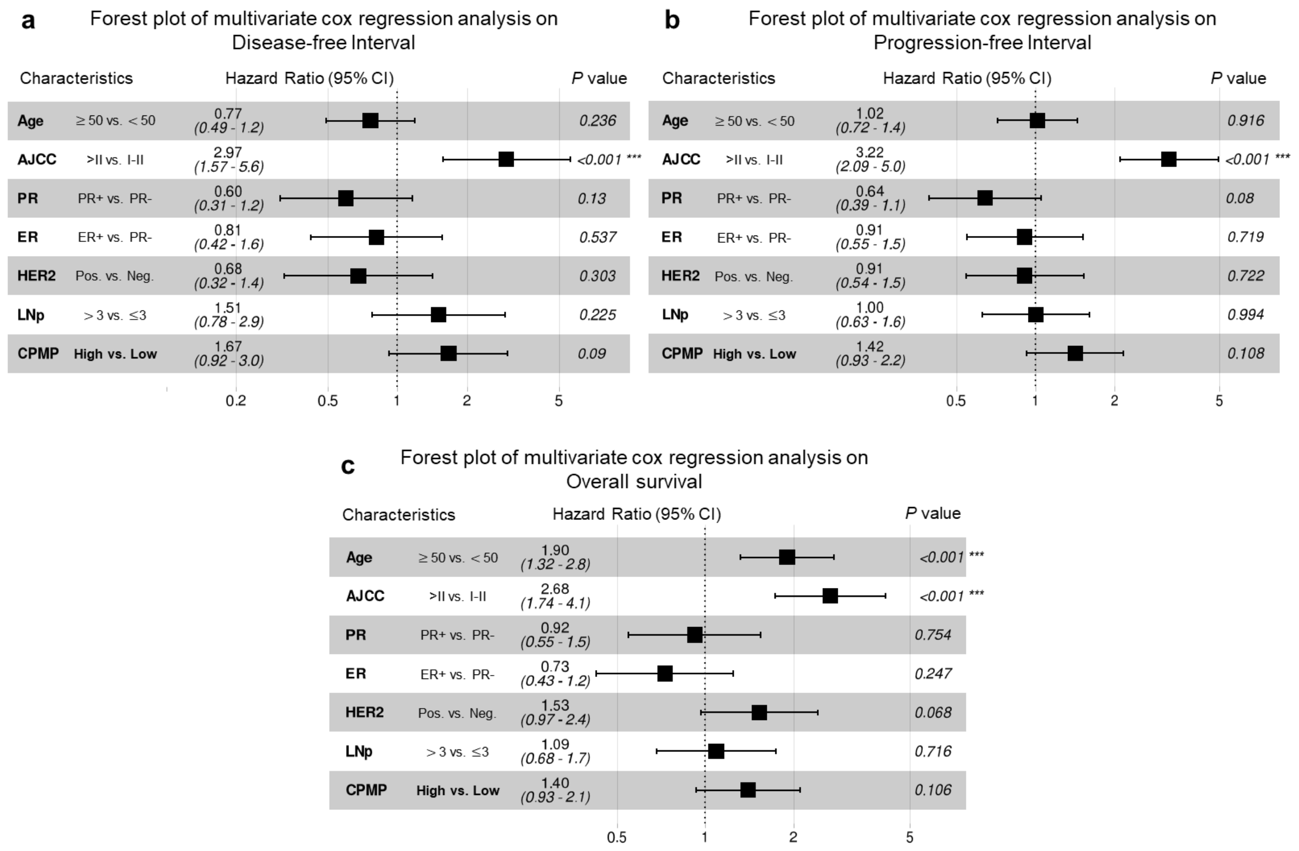


**Figure S19. Multivariate regression analysis of DFI, PFI, and OS prognostic indicators on external TCGA-BRCA data.**

Forest plot of multivariate cox regression analysis including the covariates age, AJCC pathologic tumor stage, PR status, ER status, HER2 status, LNp, and CPMP risk group and their association with Disease-free survival (**a**), Progression-free Interval (**b**), and Overall survival (**c**). HR and 95% CI were calculated by the Cox proportional hazard model. P-value was calculated using the two-sided log-rank test. (* indicate statistical significance, * p<0.05, ** p<0.01). HR=Hazard Ratio, CI=Confidence Intervals, AJCC=The American Joint Committee on Cancer, PR=Progesterone Receptor, ER=Estrogen Receptor, HER2=Human Epidermal growth factor Receptor 2, LNp=The number of Lymph Nodes positive, HR+ =Hormone Receptor positive, LN0=Lymph Nodes negative, LN1=The number of Lymph Nodes positive≤3.

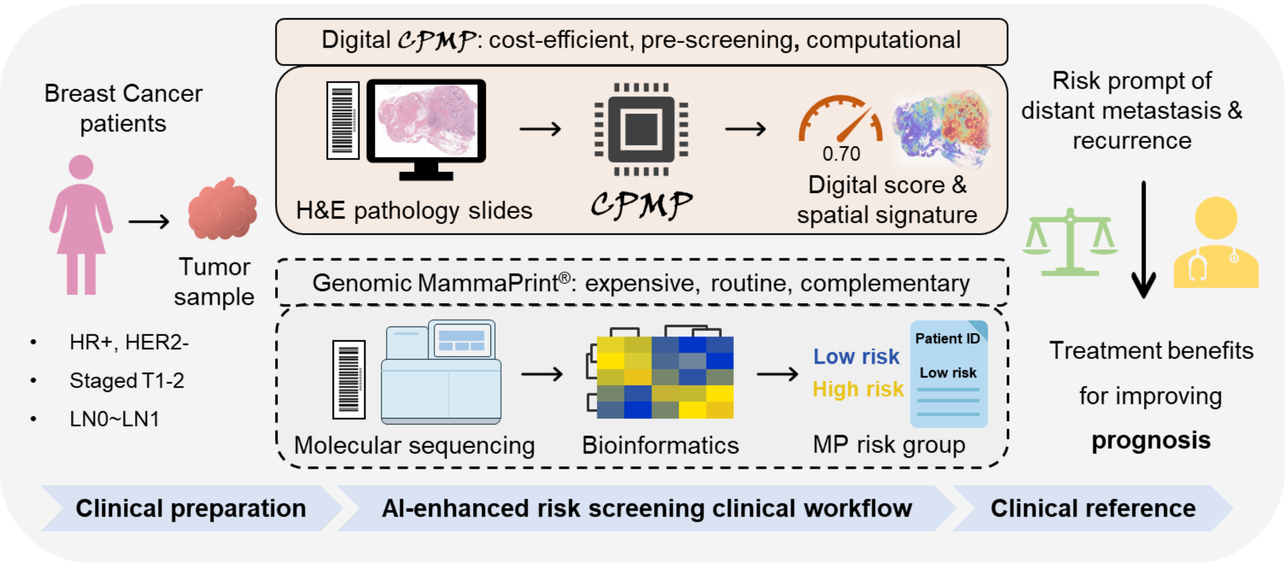


**Figure S20. Summary of digital CPMP for potential risk pre-screening in clinical workflow.**

The clinical diagnosis process could be enhanced by our proposed AI-driven computational pathology framework. CPMP provides valuable insights into the close correlations between molecular gene activities, spatial morphology in the tumor ecosystem, and prognostic knowledge. It has the potential to be a pre-screening and cost-efficient diagnostic tool for recurrence risk assessment in early-stage breast cancer patients, enabling a precise therapeutic regimen at the clinical reference phase for prognosis improvement. HR+ =Hormone Receptor positive, HER2- =Human Epidermal growth factor Receptor 2 negative, LN0=Lymph Nodes negative, LN1=The number of Lymph Nodes positive $\leq 3$, MP=MammaPrint diagnostic test.

**TableS1. Patient characteristics of the TJMUCH-MP clinical cohort.**

| Clinical variables | Total (N = 477) | High (N = 178) | Low (N = 299) | *P* value |
|---|---|---|---|---|
| Age | 52.7 (10.5) | 52.4 (10.8) | 52.8 (10.3) | 0.714 |
| Menopausal Status | | | | 0.961 |
| No / Unknown | 203 (42.6%) | 75 (42.1%) | 128 (42.8%) | |
| Yes | 274 (57.4%) | 103 (57.9%) | 171 (57.2%) | |
| Fertility Status | | | | 0.026 |
| No / Unkown | 19 (3.98%) | 2 (1.12%) | 17 (5.69%) | |
| Yes | 458 (96.0%) | 176 (98.9%) | 282 (94.3%) | |
| No. of pregnancies | 2.39 (1.19) | 2.39 (1.00) | 2.39 (1.29) | 0.947 |
| No. of birth | 1.29 (0.59) | 1.38 (0.55) | 1.24 (0.61) | 0.013 |
| No. of abortions | 1.06 (1.06) | 1.01 (1.04) | 1.08 (1.08) | 0.469 |
| Breast Feeding | | | | 0.052 |
| No / Unknown | 31 (6.50%) | 6 (3.37%) | 25 (8.36%) | |
| Yes | 446 (93.5%) | 172 (96.6%) | 274 (91.6%) | |
| History of benign breast tumor | | | | 0.672 |
| No / Unknown | 449 (94.1%) | 166 (93.3%) | 283 (94.6%) | |
| Yes | 28 (5.87%) | 12 (6.74%) | 16 (5.35%) | |
| History of malignant tumors | | | | 1 |
| No / Unknown | 460 (96.4%) | 172 (96.6%) | 288 (96.3%) | |
| Yes | 17 (3.56%) | 6 (3.37%) | 11 (3.68%) | |
| Family history of malignant tumors | | | | 0.23 |
| No / Unknown | 359 (75.3%) | 128 (71.9%) | 231 (77.3%) | |
| Yes | 118 (24.7%) | 50 (28.1%) | 68 (22.7%) | |
| Family history of breast cancer | | | | 0.458 |
| No / Unknown | 446 (93.5%) | 164 (92.1%) | 282 (94.3%) | |
| Yes | 31 (6.50%) | 14 (7.87%) | 17 (5.69%) | |
| Tumor Location | | | | 0.717 |
| Left | 253 (53.0%) | 92 (51.7%) | 161 (53.8%) | |
| Right | 224 (47.0%) | 86 (48.3%) | 138 (46.2%) | |
| Surgery Option | | | | 0.244 |
| full-cut | 239 (50.1%) | 91 (51.1%) | 148 (49.5%) | |
| Breast-conserving | 118 (24.7%) | 37 (20.8%) | 81 (27.1%) | |
| Breast-reconstruction | 120 (25.2%) | 50 (28.1%) | 70 (23.4%) | |
| T | | | | 0.412 |
| T1 | 294 (61.6%) | 105 (59.0%) | 189 (63.2%) | |
| T2 | 183 (38.4%) | 73 (41.0%) | 110 (36.8%) | |
| N | | | | 0.002 |
| N0 | 362 (75.9%) | 137 (77.0%) | 225 (75.3%) | |
| 1mic | 35 (7.34%) | 21 (11.8%) | 14 (4.68%) | |
| 1a | 80 (16.8%) | 20 (11.2%) | 60 (20.1%) | |
| AJCC | | | | <0.001 |
| IA | 252 (52.8%) | 83 (46.6%) | 169 (56.5%) | |
| IB | 22 (4.61%) | 19 (10.7%) | 3 (1.00%) | |
| IIA | 142 (29.8%) | 61 (34.3%) | 81 (27.1%) | |
| IIB | 61 (12.8%) | 15 (8.43%) | 46 (15.4%) | |
| Histological grading | | | | <0.001 |
| I | 14 (2.94%) | 5 (2.81%) | 9 (3.01%) | |
| I~II | 45 (9.43%) | 9 (5.06%) | 36 (12.0%) | |
| II | 393 (82.4%) | 144 (80.9%) | 249 (83.3%) | |
| II~III | 15 (3.14%) | 12 (6.74%) | 3 (1.00%) | |
| III | 10 (2.10%) | 8 (4.49%) | 2 (0.67%) | |
| Lymphatic Invasion | | | | 0.968 |
| No | 423 (88.7%) | 159 (89.3%) | 264 (88.3%) | |
| Yes | 43 (9.01%) | 15 (8.43%) | 28 (9.36%) | |
| suspicious | 11 (2.31%) | 4 (2.25%) | 7 (2.34%) | |
| TILs | 3.36 (8.32) | 5.34 (11.5) | 2.19 (5.29) | 0.001 |
| Total number of axillary lymph nodes removed | 10.1 (7.77) | 11.2 (8.49) | 9.49 (7.24) | 0.028 |
| Number of positive axillary lymph nodes | 0.20 (0.47) | 0.13 (0.38) | 0.24 (0.51) | 0.009 |
| LN_ratio | 0.02 (0.05) | 0.01 (0.03) | 0.02 (0.05) | <0.001 |
| ER | 86.2 (10.4) | 85.5 (11.7) | 86.7 (9.52) | 0.236 |
| PR | 61.9 (31.7) | 52.8 (34.3) | 67.4 (28.7) | <0.001 |
| HER2 | | | | 0.081 |
| HER2-negtaive | 139 (29.1%) | 43 (24.2%) | 96 (32.1%) | |
| HER2-low | 338 (70.9%) | 135 (75.8%) | 203 (67.9%) | |
| Ki67 | 20.8 (12.4) | 27.8 (14.6) | 16.6 (8.57) | <0.001 |

**TableS2. Evaluation performance of CPMP and comparative methods for MP recurrence risk prediction.**

| Performance | AUROC | Balanced accuracy | Weighted avg. recall | Weighted avg. precision | Weighted avg. F1-score | High-risk F1-score | Low-risk F1-score |
|---|---|---|---|---|---|---|---|
| **CPMP** | **0.824±0.029** | **0.772±0.030** | **0.777±0.027** | **0.786±0.027** | **0.778±0.027** | **0.715±0.036** | **0.816±0.026** |
| CEMIL | 0.741±0.039 | 0.717±0.030 | 0.737±0.031 | 0.742±0.028 | 0.734±0.030 | 0.643±0.047 | 0.789±0.033 |
| CLAM | 0.798±0.044 | 0.752±0.044 | 0.765±0.042 | 0.770±0.039 | 0.764±0.043 | 0.690±0.059 | 0.809±0.040 |
| TransMIL | 0.755±0.056 | 0.711±0.044 | 0.718±0.053 | 0.737±0.045 | 0.717±0.052 | 0.644±0.052 | 0.760±0.065 |
| Wagner et al. | 0.806±0.027 | 0.759±0.024 | 0.764±0.032 | 0.776±0.024 | 0.765±0.030 | 0.701±0.029 | 0.803±0.037 |
| ABMIL | 0.810±0.028 | 0.759±0.036 | 0.762±0.040 | 0.776±0.033 | 0.763±0.039 | 0.701±0.043 | 0.801±0.043 |
| DTFDMIL | 0.819±0.027 | 0.768±0.028 | 0.770±0.037 | 0.783±0.026 | 0.771±0.036 | 0.713±0.030 | 0.806±0.043 |
| SAMMIL | 0.770±0.036 | 0.719±0.033 | 0.726±0.046 | 0.749±0.030 | 0.724±0.045 | 0.652±0.046 | 0.767±0.064 |

Mean and standard deviation are reported, and the best results are marked in **bold**.
MP=MammaPrint diagnostic test
AUROC=Area Under the Receiver Operating Characteristic

**TableS3. Evaluation performance of the models in filtering noisy tissue.**

| Performance | Evaluation metrics | | | | | |
|---|---|---|---|---|---|---|
| | **High-risk recall** | **Low-risk recall** | **Weighted avg. F1-score** | **Balanced accuracy** | **AUROC** | ***p*-value** |
| w/o noisy tissue filtering (CPMP) | 0.750±0.075 | 0.793±0.053 | 0.778±0.027 | 0.772±0.030 | 0.824±0.029 | ns |
| w/ noisy tissue filtering | 0.764±0.071 | 0.779±0.067 | 0.775±0.034 | 0.772±0.033 | 0.826±0.032 | |

Mean and standard deviation are reported.
w/=with, w/o=without, ns=Statistically no significance.
AUROC=Area Under the Receiver Operating Characteristic

**TableS4. Evaluated AUROC values depending on the number of tiles sampled in each slide.**

| **AUROC values** | | **The sampling number of tiles / each slide** | | | | | | | | | |
|---|---|---|---|---|---|---|---|---|---|---|---|
| | | 1 | 5 | 10 | 50 | 100 | 500 | 1000 | 5000 | 10000 | All |
| **Trained on sampling size** | 1k | 0.63 | - | 0.76 | - | 0.82 | - | 0.824 | - | 0.824 | 0.82 |
| | 5k | 0.63 | 0.72 | 0.76 | 0.81 | 0.82 | 0.82 | 0.825 | 0.82 | 0.824 | 0.82 |
| | 10k | 0.62 | 0.71 | 0.76 | 0.81 | 0.81 | 0.82 | 0.824 | 0.82 | 0.823 | 0.82 |
| | all | 0.62 | 0.71 | 0.75 | 0.8 | 0.81 | 0.82 | 0.823 | 0.82 | 0.824 | 0.82 |

Mean value is reported for each experiment.
AUROC=Area Under the Receiver Operating Characteristic

**TableS5. Clinicopathological variable mapping table for the clinical-based model development.**

| **Clinical variables** | **Numerical codes** | | | | |
|---|---|---|---|---|---|
| | 0 | 1 | 2 | 3 | 4 |
| Surgery Option | Full-cut | Breast-conserving | Breast-reconstruction | - | - |
| T stage | - | T1 | T2 | - | - |
| N stage | N0 | 1mic | 1a | 2a | - |
| AJCC | IA | IB | IIA | IIB | IIIA |
| Histological grading | I | I~II | II | II~III | III |

**TableS6. Evaluation performance of our CPMP and its comparison with the clinical-based model.**

| Performance | Evaluation metrics | | | | | | |
|---|---|---|---|---|---|---|---|
| | AUROC | Balanced accuracy | Weighted avg. recall | Weighted avg. precision | Weighted avg. F1-score | High-risk F1-score | Low-risk F1-score |
| CPMP | **0.824±0.029** | **0.772±0.030** | **0.777±0.027** | **0.786±0.027** | **0.778±0.027** | **0.715±0.036** | **0.816±0.026** |
| Clinical-based | 0.800±0.041 | 0.698±0.026 | 0.748±0.022 | 0.748±0.025 | 0.734±0.024 | 0.596±0.042 | 0.816±0.016 |

Mean and standard deviation are reported, and the best results are marked in **bold**.
AUROC=Area Under the Receiver Operating Characteristic

**TableS7. Evaluation performance of agent-transformer models with variant number of the agent token.**

| The number of the agent token | Evaluation metrics | | | | | | |
|---|---|---|---|---|---|---|---|
| | AUROC | Balanced accuracy | Weighted avg. recall | Weighted avg. precision | Weighted avg. F1-score | High-risk F1-score | Low-risk F1-score |
| 1 | 0.824±0.029 | 0.772±0.030 | 0.777±0.027 | 0.786±0.027 | 0.778±0.027 | 0.715±0.036 | 0.816±0.026 |
| 2 | 0.822±0.035 | 0.772±0.032 | 0.777±0.034 | 0.788±0.029 | 0.777±0.033 | 0.716±0.038 | 0.814±0.036 |
| 4 | 0.816±0.037 | 0.766±0.035 | 0.772±0.034 | 0.783±0.031 | 0.772±0.033 | 0.709±0.043 | 0.811±0.035 |
| 8 | 0.817±0.034 | 0.763±0.032 | 0.778±0.028 | 0.783±0.029 | 0.777±0.027 | 0.702±0.040 | 0.822±0.026 |
| 16 | 0.811±0.032 | 0.764±0.032 | 0.767±0.034 | 0.780±0.028 | 0.768±0.034 | 0.706±0.038 | 0.804±0.039 |
| 32 | 0.815±0.031 | 0.768±0.031 | 0.777±0.038 | 0.788±0.028 | 0.776±0.036 | 0.711±0.037 | 0.815±0.044 |
| 64 | 0.810±0.036 | 0.760±0.035 | 0.769±0.030 | 0.777±0.029 | 0.769±0.030 | 0.699±0.047 | 0.811±0.028 |
| 128 | 0.800±0.033 | 0.747±0.028 | 0.758±0.036 | 0.768±0.027 | 0.757±0.034 | 0.685±0.036 | 0.801±0.042 |
| 256 | 0.802±0.032 | 0.756±0.026 | 0.765±0.031 | 0.775±0.025 | 0.765±0.029 | 0.696±0.032 | 0.807±0.035 |
| Self-attention Transformer | 0.806±0.027 | 0.759±0.024 | 0.764±0.032 | 0.776±0.024 | 0.765±0.030 | 0.701±0.029 | 0.803±0.037 |
| **CPMP** | **0.824±0.029** | **0.772±0.030** | **0.777±0.027** | **0.786±0.027** | **0.778±0.027** | **0.715±0.036** | **0.816±0.026** |

Mean and standard deviation are reported, and the best results are marked in **bold**.
AUROC=Area Under the Receiver Operating Characteristic

**TableS8. Evaluation performance of the models equipped with different positional encoding configurations.**

| Positional encoding configurations | Evaluation metrics | | | | |
|---|---|---|---|---|---|
| | AUROC | Balanced accuracy | Weighted avg. recall | Weighted avg. precision | Weighted avg. F1-score |
| PE | **0.824±0.029** | 0.772±0.030 | **0.777±0.027** | 0.786±0.027 | **0.778±0.027** |
| no PE | 0.820±0.031 | **0.773±0.030** | 0.768±0.032 | 0.788±0.027 | 0.770±0.031 |
| PPEG | 0.821±0.026 | 0.771±0.028 | 0.772±0.035 | **0.788±0.023** | 0.773±0.034 |

Mean and standard deviation are reported, and the best results are marked in **bold**.
AUROC=Area Under the Receiver Operating Characteristic
PE=Positional coordinates information Embeddings, PPEG=Pyramid Position Encoding Generator

**TableS9. Evaluation performance of the experimental models equipped with different loss functions.**

| Different loss functions | Evaluation metrics | | | | |
|---|---|---|---|---|---|
| | AUROC | Balanced accuracy | Weighted avg. recall | Weighted avg. precision | Weighted avg. F1-score |
| MSE+MSE | **0.824±0.029** | **0.772±0.030** | **0.777±0.027** | **0.786±0.027** | **0.778±0.027** |
| MSE+CE | 0.820±0.031 | 0.766±0.031 | 0.770±0.031 | 0.782±0.027 | 0.770±0.031 |
| MSE | 0.820±0.031 | 0.766±0.032 | 0.769±0.031 | 0.783±0.030 | 0.770±0.030 |
| CE | 0.812±0.028 | 0.762±0.024 | 0.767±0.023 | 0.778±0.021 | 0.768±0.022 |

Mean and standard deviation are reported, and the best results are marked in **bold**.
AUROC=Area Under the Receiver Operating Characteristic
MSE=Mean Square Error, CE=Cross Entropy

**TableS10. Evaluation performance of the models equipped with different feature extraction configurations.**

| Different feature extraction configurations | Evaluation metrics | | | | | | |
|---|---|---|---|---|---|---|---|
| | Model Params. | Dim. Of Embed. | AUROC | Balanced accuracy | Weighted avg. recall | Weighted avg. precision | Weighted avg. F1-score |
| UNI | 307 M | 1024 | 0.824±0.029 | 0.772±0.030 | 0.777±0.027 | 0.786±0.027 | 0.778±0.027 |
| Prov-GigaPath | 1.1 B | 1536 | 0.827±0.022 | 0.772±0.028 | 0.773±0.036 | 0.790±0.024 | 0.774±0.035 |
| Virchow | 632 M | 2560 | 0.827±0.033 | 0.770±0.030 | 0.782±0.030 | 0.791±0.022 | 0.781±0.029 |
| H-optimus-0 | 1.1 B | 1536 | 0.824±0.029 | 0.779±0.026 | 0.788±0.032 | 0.797±0.025 | 0.788±0.031 |
| Phikon | 85.8 M | 768 | 0.804±0.037 | 0.755±0.035 | 0.759±0.036 | 0.774±0.029 | 0.759±0.036 |
| CTransPath | 28.3 M | 768 | 0.740±0.031 | 0.701±0.026 | 0.712±0.038 | 0.727±0.025 | 0.711±0.035 |

Mean and standard deviation are reported.
AUROC=Area Under the Receiver Operating Characteristic
B=Billion, M=Million

**TableS11. Slide-level performance metrics of spatial attention maps compared with pathologist-labeled tumor regions across 20 WSIs**

| SlideID | Evaluation metrics | | | | |
|---|---|---|---|---|---|
| | **AUC** | **Recall Ratio** | **Overlap Ratio** | **IoU** | **DICE** |
| 5067 | 0.890 | 0.512 | 0.744 | 0.435 | 0.606 |
| 1060 | 0.941 | 0.613 | 0.432 | 0.339 | 0.507 |
| 1052 | 0.873 | 0.271 | 0.968 | 0.268 | 0.423 |
| 1081 | 0.925 | 0.545 | 0.794 | 0.477 | 0.646 |
| 1068 | 0.964 | 0.760 | 0.550 | 0.469 | 0.638 |
| 2042 | 0.982 | 0.825 | 0.445 | 0.407 | 0.578 |
| 4020 | 0.978 | 0.643 | 0.925 | 0.611 | 0.759 |
| 1057 | 0.941 | 0.667 | 0.788 | 0.566 | 0.723 |
| 1042 | 0.853 | 0.291 | 0.933 | 0.285 | 0.444 |
| 1013 | 0.919 | 0.512 | 0.833 | 0.464 | 0.634 |
| 1094 | 0.944 | 0.685 | 0.673 | 0.514 | 0.679 |
| 2055 | 0.955 | 0.710 | 0.731 | 0.563 | 0.720 |
| 2050 | 0.959 | 0.622 | 0.479 | 0.371 | 0.541 |
| 2010 | 0.920 | 0.410 | 0.826 | 0.378 | 0.548 |
| 1051 | 0.970 | 0.660 | 0.765 | 0.549 | 0.709 |
| 1066 | 0.996 | 0.717 | 0.578 | 0.471 | 0.640 |
| 5004 | 0.961 | 0.618 | 0.906 | 0.581 | 0.735 |
| 3090 | 0.990 | 0.883 | 0.212 | 0.206 | 0.342 |
| 3046 | 0.970 | 0.755 | 0.693 | 0.565 | 0.722 |
| 3017 | 0.993 | 0.902 | 0.796 | 0.733 | 0.846 |
| **Mean** | **0.946** | **0.630** | **0.704** | **0.463** | **0.622** |
| Std. | 0.040 | 0.171 | 0.198 | 0.130 | 0.126 |

Evaluation performance is reported for each WSI, mean (marked in **bold**) and standard deviation across the 20 WSIs are shown in the last two rows.
AUC = area under the receiver operating characteristic curve.
Recall Ratio=intersection over union between CPMP-generated attention regions and pathologist-labeled tumor annotations relative to pathologist-labeled tumor annotations.
Overlap Ratio=intersection over union between CPMP-generated attention regions and pathologist-labeled tumor annotations relative to CPMP-generated attention regions.
IoU = intersection over union.
DICE = Dice similarity coefficient.
